# A simple cognitive model explains movement decisions during schooling in zebrafish

**DOI:** 10.1101/2023.02.05.527161

**Authors:** Lital Oscar, Liang Li, Dan Gorbonos, Iain D. Couzin, Nir S. Gov

## Abstract

While moving, animals must frequently make decisions about their future travel direction, whether they are alone or in a group. Here we investigate this process for zebrafish (*Danio rerio*), which naturally move in cohesive groups. Employing state-of-the-art virtual reality, we study how real fish follow one or several moving, virtual conspecifics. These data are used to inform, and test, a model of social response that includes a process of explicit decision-making, whereby the fish can decide which of the virtual conspecifics to follow, or to follow some average direction. This approach is in contrast with previous models where the direction of motion was based on a continuous computation, such as directional averaging. Building upon a simplified version of this model [Sridhar et al., 2021], which has been shown to exhibit a spontaneous symmetry-breaking transition from moving along a “compromise” (average) direction, to deciding on following one of the virtual fish. This previously published simplified version was limited to a one-dimensional projection of the fish motion, while here we present a model that describes the motion of the real fish as it swims freely in two-dimensions. Here, we extend our proposed Ising-like model, which inherently describes a spontaneous symmetry-breaking transition from moving along a “compromise” (average) direction, to deciding on following one of the virtual fish. Motivated by experimental observations, the swim speed of the fish in this model uses a burst-and-coast swimming pattern, with the burst frequency being dependent on the distance of the fish from the followed conspecific(s). We demonstrate that this model is able to explain the observed spatial distribution of the real fish behind the virtual conspecifics in the experiments, as a function of their average speed and number. In particular, the model naturally explains the observed critical bifurcations for a freely swimming fish, which appear in the spatial distributions whenever the fish makes a decision to follow only one of the virtual conspecifics, instead of following them as an averaged group. This model can provide the foundation for modeling a cohesive shoal of swimming fish, while explicitly describing their directional decision-making process at the individual level.

## I. INTRODUCTION

Understanding how animals move together in groups is a long-standing puzzle [1, 2]. Current theoretical models of this process involve the description of individual agents that move according to simple rules, which embody the social influence exerted by their conspecifics, and external influences such as physical barriers and predators [2–8]. Within these models, the influences of the neighbors in the group, representing the sensory information, are integrated, and the resulting direction of motion of each agent is calculated. However, during this calculation, the individuals in such model frameworks do not explicitly perform a decision-making process. In other words, the direction of motion of each agent is given by some unique and continuous function of the positions and velocities of the detected (usually neighboring) conspecifics [3, 4, 9–13]. Given two identical conspecifics, the agent in these modelas has no mechanism to spontaneously decide to follow only one of them (at a given time), in contradiction to the observations [14].

We propose here a theoretical model for how animals respond to social cues, that includes an explicit decision-making process [14]. Our current work is based on our theoretical model for how animal groups, and individual animals, make decisions regarding their desired direction of motion while moving towards and choosing between different targets [14, 15]. This assumption is supported by neurobiological studies conducted in a range of animals [16, 17]. Within this model, the animal’s brain is assumed to have neuronal representations of the vectors aimed at the different targets. The neurons (or neuronal groups) that represent each target are treated as “spins”, in the terminology of statistical physics, with each spin representing either an “on” (1) or “off” (0) state. The spin state corresponds to the firing state of the neuronal group that it represents. This spin model is a simplification of (and can be mapped to [14]) widely used models of neuronal representation of spatial targets [18, 19]. The interactions between the spins drives a geometry-based phase transition (from averaging among options to deciding on one), which manifest as bifurcations in the resulting trajectories. This model thus predicts that the animal’s direction of travel would tend to be in a compromise (average) direction between the different targets, when they are far, and the relative angle between them is small. However, above a critical relative angle, such as when the animal approaches the targets, the trajectories bifurcate and point toward one target, in the case of only 2 options, and a subset of targets for a larger number of options. This prediction was verified for several types of insects (desert locusts and fruit flies), moving toward stationary targets in a virtual-reality arena [14].

The model was also employed to consider how zebrafish (*Danio rerio*) respond to conspecifics. Again using immersive virtual reality [14], allowing virtual fish to be projected in specific configurations while on the move, it was also found that individuals would average when the angle subtended by virtual fish was below a critical value, and a decision to follow one among the remaining options occurred when this angle exceeded a critical value. In the comparison with the theoretical model in this previous work [14], it was assumed that the real fish (RF) can only move along one dimension, perpendicular to the direction of motion of the virtual fish (VF) (Fig. 1). This way, the RF is assumed to be moving at a fixed distance behind the line of VF, and only its lateral motion sideways was modeled. Note that in the experiments, the RF also maintains a very similar depth to the VF that it is chasing [14], thus schools of real juvenile zebrafish tend to be quasi-planar. Despite the considerable simplification in the theoretical model, the experimental and simulated distributions of the RF, when projected to one dimension, were in very good agreement for both two and three VF [14]. The essence of the simplification obtained by the one-dimensional projection of the fish motion is that it eliminates the need to calculate the speed of the fish, as it responds to the VF.

**FIG. 1:**
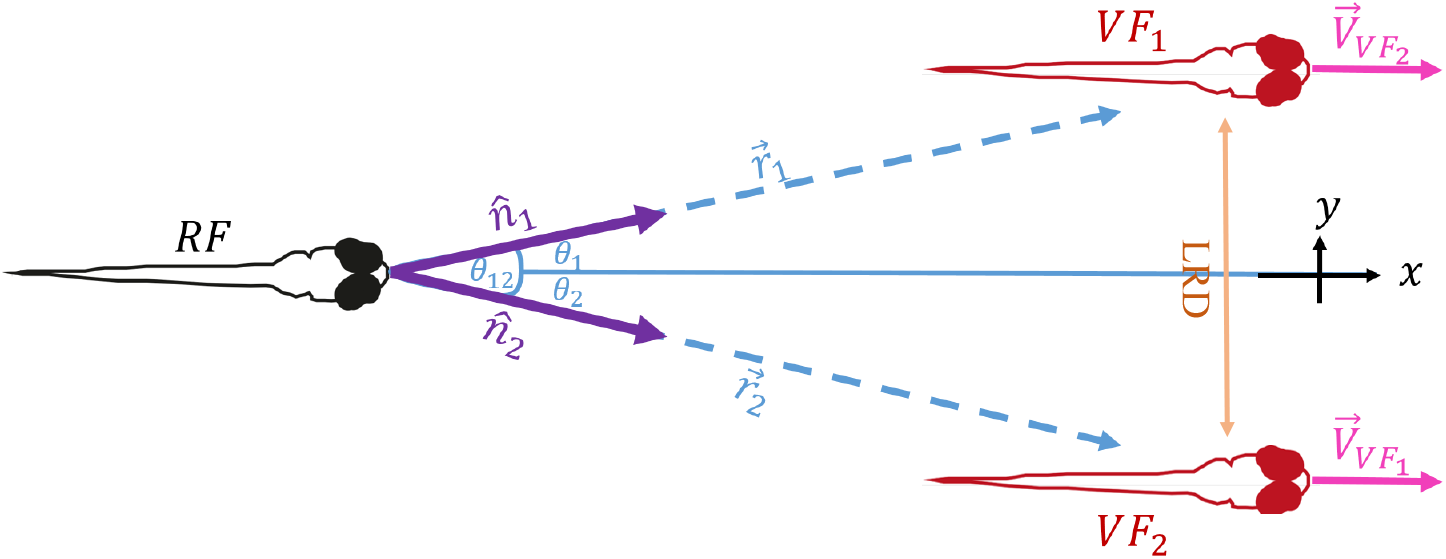
Illustration of a 2D system with 1RF (black) following 2VF (red): *θ*_12_ is the instantaneous relative angle between the two targets, *θ*_*i*_ is the angle from the RF to *V F*_*i*_. Note that we use the same terminology to describe our model as employed in our experiments; thus, the RF in the model represents the focal individual whose cognitive process is being simulated whereas the VF represent the VF used in the virtual-reality experiments, moving in the same direction as each other and at a fixed distance apart. The instantaneous direction vectors to the targets are 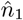 and 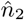, with the corresponding distances *r*_1_,*r*_2_. The lateral distance (LRD) denotes the distance between nearest neighbor VF, when the VF are moving along a line that is perpendicular to their direction of motion (modified from [14]). The VF velocity vectors were taken to be identical, so that the VF move in synchrony, and describes a series of burst-and-coast dynamics.

In the experiments, however, the RF moves freely in (predominantly) two dimensions (due to the nature of fish to swim at the same depth as each other) when schooling with the VF, and its distance to the VF constantly changes during the chase. Here we take this freedom of movement into account and develop a theoretical framework that implements the spin model for directional decision-making, together with a calculation of the fish speed, as the RF interacts freely with an arbitrary number and spatial configurations of moving conspecifics (which can be thought of as mobile targets). This model allows us for the first time to compare the spin model for directional decision-making to the full two-dimensional experimental trajectories of the RF [14]. We emphasize that our model does not include any explicit alignment interaction of the Vicsek-type [2, 20]. Moreover, the main theoretical novelty of our work is the spin-based directional decision making model, with its spontaneous symmetry-breaking property, as opposed to previous models using vectorial averaging and simple distance-based interaction rules. The calculation of the speed, using burst-and-coast dynamics, is more standard [21, 22].

## II. THE FISH MODEL

Motivated by our experimental observations on zebrafish (*Danio rerio*) [14], we will limit the motion of the real fish (RF) in our model to a two-dimensional plane on which fish interact (i.e. they are assumed to be swimming at the same depth in the water column) (Fig. 1). In our experiments the VF move with the burst-and-coast (or, burst-and-glide) motions as do real fish (Fig. 2A). In these experiments the virtual fish move in straight trajectories, and make U-turns when they near the edge of the experimental arena. Our model allows us to calculate the full two dimensional motion of the RF, when it interacts with several VF (which move as in the experiments), as illustrated in Fig. 1. We clearly denote in all the figures the experimental data versus the theoretical results, to allow a direct visual comparison to be made.

**FIG. 2:**
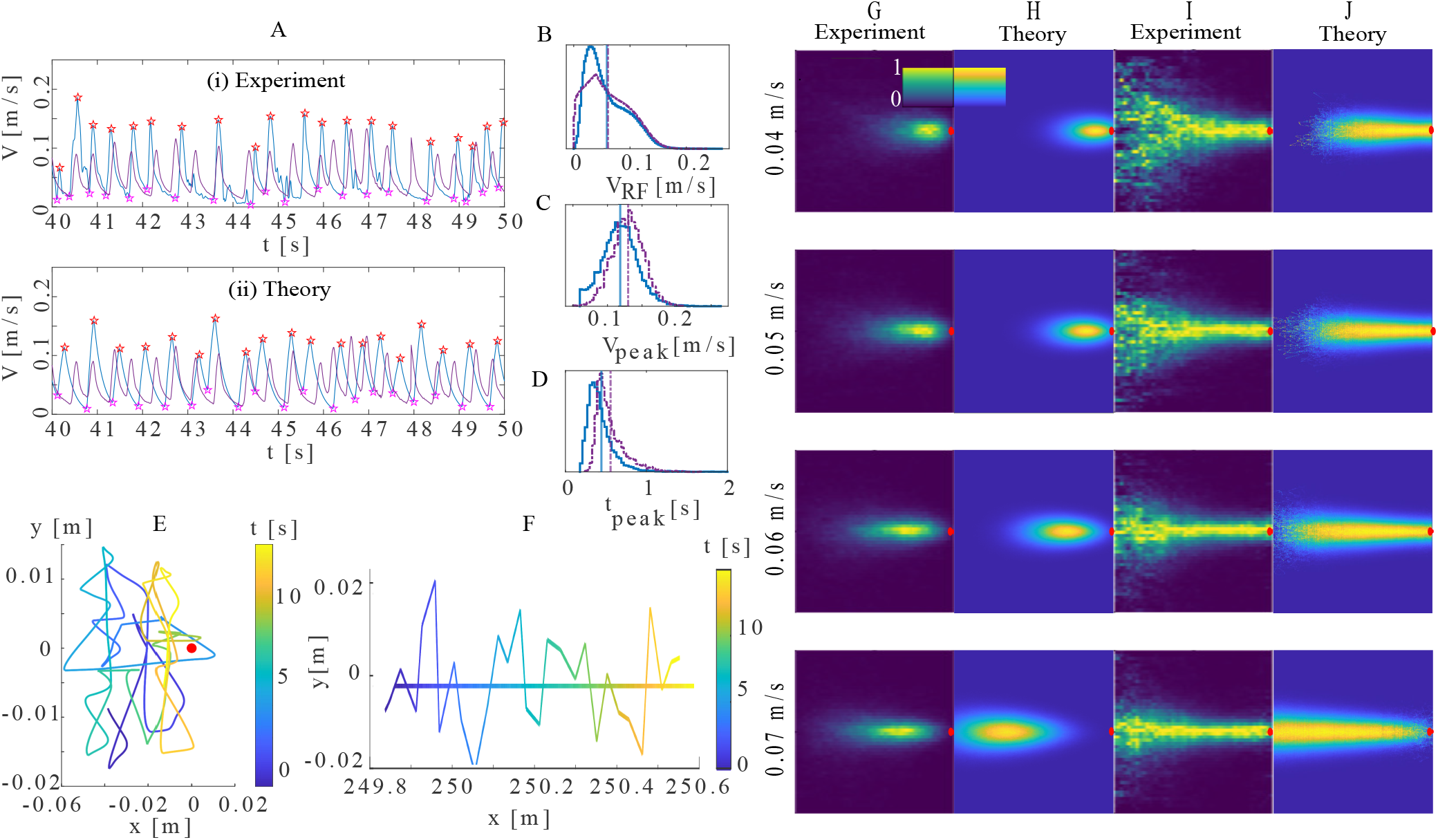
RF following one VF: burst-and-coast dynamics and spatial distributions. (A) (i) The RF’s speed dynamics in the experiment (blue) compared with the VF velocity (purple, with an average velocity of *V*_*V F*_ = 0.05[*m/s*]), and (ii) shows the simulated RF velocity (blue) compared with the VF velocity (purple). The maxima and minima of the speed are denoted by the red and pink stars respectively. (B) The speed distribution of the RF, (C) The distribution of the RF’s velocity peak values, and (D) The distribution of time intervals between consecutive bursts (*t*_*peak*_). In (B-D) the purple line denotes the simulation and blue line the experimental data. (E) An example of the RF trajectory behind 1VF, relative to the VF movement (located at the red dot). (F) Same as (E) but in the lab frame, where the horizontal line gives the trajectory of the VF. In both (E) and (F) time is represented by the color bar. (G-J) Accumulated distribution of the RF behind the VF, for different mean *V*_*V F*_ (each row), in the VF frame of reference (the VF is at the origin, red dot). For each velocity, we normalized the heat maps over the whole 2D space (G and H), or over individual *x*-sections (I and J). (G) and (I) show the experimental heat maps, (H) and (J) the simulated results. For each VF velocity, we ran 100 simulations (in which the RF initial position was random, within a certain distance of the VFs) each for 5, 000[*s*] (500,000 iteration steps). We used the following parameters: *T* = 0.1, *η* = 5[*s*^−1^], *γ* = 5[*s*^−1^], *σ* = *π/*3[*rad/s*] (standard deviation of the angular noise), *b* = *±π*[*rad/s*] (angular noise limits), *k* = 250[(*m s*)^−1^], *r*_*d*_ = 0.2[*m*], *t*_*off*_ = 0.15[*s*], *f*_0_ = 1.1[*m/s*^2^].

### A. Direction of motion: Spin dynamics

Within our model the direction in which the RF is moving is dictated by the neuronal firing, which is represented by the “on” spins in the Ising-like model [14]. Each target is described by a group of spins; each one can be either “on” (1) or “off” (0), such that the net unit vector of the RF’s desired direction of motion is given by

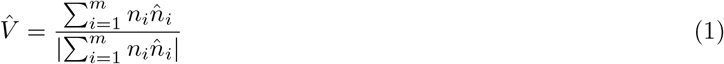

where, *n*_*i*_ refers to the number of “on” spins in subgroup *i*, 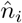 is the unit vector pointing from the RF towards target *i* (Fig. 1), and *m* is the number of VF (and spin groups).

Each neural group vector 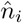 is directed, on average, at each of the different VF, but we consider that there is angular noise affecting the angular direction of this vector according to the following Langevin equation

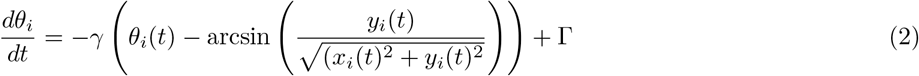

where *x*_*i*_, *y*_*i*_ are the coordinates of target *i* with respect to the location of the RF, Γis the amplitude of the Gaussian noise (with a standard deviation of 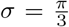 and limits of −*b* < Γ< *b*, where *b* = *π*), and *γ* is the rate at which the angle of the internal vector re-adjusts to the new perspective angle of target *i*.

The dynamics of the spins in each group are calculated using the Ising model, based on the following Hamiltonian [14, 15]

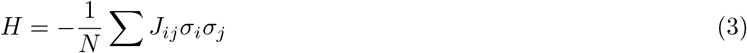

where *σ*_*i*_ are the spin variables, *N* is the total number of spins in all subgroups, and *J*_*ij*_ are the interactions between the spins, given as the dot products of the preferred directions of spins *i* and *j*, such that: 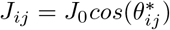, where *J*_0_ is the interaction strength energy, and 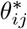 is the modified form of the relative angle according to:

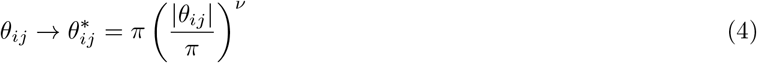

where *θ*_*ij*_ is the Euclidean relative angle (Fig. 1) and the tuning parameter *ν* describes the non-Euclidean transformation of the relative angle, thereby determining the interactions between the spins (Fig. S1). Comparisons with experiments indicate that a suitable tuning parameter is *ν* = 0.5 [14], across taxa, which is the value we used in our simulations. The dynamics of the spins follow from the Hamiltonian (Eq. 3). They can be calculated for discrete, stochastic spins, or using a continuum description in terms of the following Langevin equation for the dynamics of the neural firing

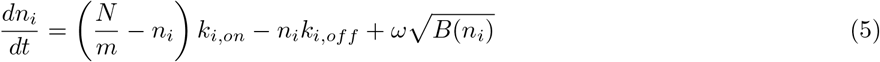

where *B*(*n*_*i*_) is the noise amplitude, *m* is the number of neuronal groups, and *ω* is a normally distributed white Gaussian noise (with a standard deviation of 1). This dynamic is applied to each spin group that represents a target. The noise amplitude *B*(*n*_*i*_) is given by [15] (see SI section S2)

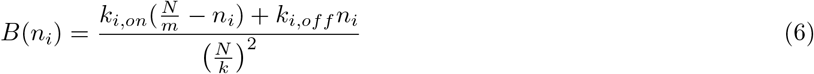

where the spin-flip rates (per spin) are given by

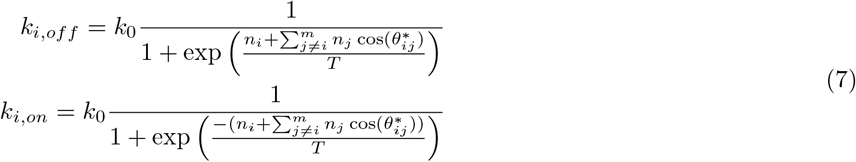

where the sum goes over all spin groups except *i, k*_0_ = 1[*s*^−1^] is a spin-flip rate constant, and *T* stands for the dimensionless ratio between the noise amplitude and the spin-spin interaction strength, which determines the dynamics of the spins flips. These spin flips represent the dynamics of neuronal switching between firing and non-firing states. Integrating Eq. 5 over time gives us the dynamics of the spins that are “on”, which determines the direction of the RF motion (Eq. 1), as a function of time.

The contribution of each spin group to the desired direction in Eq. 1 is modified if there are several targets that are very close to each other. Previous work suggests that targets that are within a small relative angle with respect to the animal, are considered as a single target [23]. To account for this effect, we incorporate an ‘overlap’ function in our model [14]. This function reduces the weight of spins encoding a target in Eq. 1, if there are other targets in directional proximity (see SI section S3). This effect becomes crucially important for the case of 3VF, as described below (Fig. 4).

Note that the distance to the target does not influence an individual’s direction in the current model: spins that represent proximal targets do not have a larger weight in Eq. 1 compared to spins that represent more distant targets. We can add such modifications in the future, as needed.

### B. Speed of motion: Burst-and-coast dynamics

In this work, we will consider two possible models for the amplitude of the velocity (*V*), and in both models, this amplitude depends on the distance (*r*) between the RF and the chased VF. Recent experiments demonstrated that the effective interactions between leader-follower have a spring-like property [3, 11, 12], whereby the follower increases its speed as its distance to the leader increases, up to a critical distance where it declines. This behavior was also measured for the RF following the VF (see Fig. S4). Note that the precise implementation of this tendency, which keeps the RF from losing the VF that it is chasing, is not the main focus of our model. The novel feature that we introduce relates to directional decision making (as described in the previous section). Therefore, there can be different implementations of the RF speed, as we discuss below.

In the SI section S5, we give the results for a model where the speed of the RF (*V*_*RF*_) is described as a continuous function of the distance between the RF and VF. However, we wish to focus here on a more realistic description of the RF speed, as arising from burst-and-coast dynamics [24, 25], in which the RF’s velocity will be given by bursts of force from the tail-beat events, occurring in a stochastic manner. The differential form of the velocity can be written as

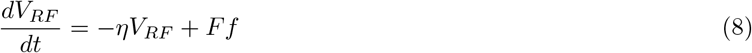

where *η* is the friction coefficient with the water, *F* is the stochastic force parameter, which is either “on” (*F* = 1) or “off” (*F* = 0), and *f* is the force amplitude of each burst (in units of force/mass). We assume that the rate at which fish exhibit bursts of tail beats (which account for the rapid acceleration in the burst-phase of their motion), *k*_*s,on*_, depends solely on the distance of the RF from its targets (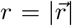, Fig. 1), and therefore we use the following expression

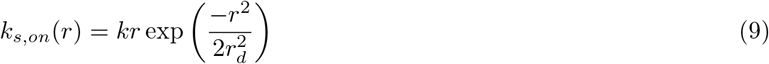

where *k* is an effective spring constant (*k* = 250[*m · s*]^−1^), and *r*_*d*_ is the length-scale beyond which the fish loses its target (we fit it to be *r*_*d*_ ∼ 0.2[*m*]). The linear dependence of the burst rate on the distance to the VF, for small values of *r* (Eq. 9), is motivated by the interactions found between fish [11], and allows the RF to successfully pursue the VF without losing it. We tested this relation by statistical analysis of the experimental RF’s dynamics, which was not conclusive (SI section S7, Fig. S6).

We use this rate in a Gillespie method (see SI section S5) within our Langevin simulations, and it determines when the next burst of tail beats occurs. Following each burst, *F* is turned “off” after a fixed duration, *t*_*off*_ = 0.15[*s*], which for simplicity was chosen to be constant in all the simulations. When the burst of tail beats stops (*F* : 1 → 0), the coasting phase starts, and the velocity of the RF decreases exponentially with time (Eq. 8), until the next burst event.

The heading direction of the simulated RF is updated at the onset of each tail burst, since in experiments it was found that the orientation of the RF stays approximately constant for the coasting duration (and is given by Eq. 1) [21, 24].

There are two additional ingredients that we found to be important to implement in our model: In order to better resemble the velocity pattern of the experimental RF, we had to define a velocity threshold (*V*_*threshold*_), whereby tail bursting occurs only when the simulated RF’s velocity is lower than this threshold (SI section S8). In Fig. S7, we compare the experimental and model RF’s velocity dynamic with and without the velocity threshold, of *V*_*threshold*_ = 0.04[*m/s*]. The velocity threshold was estimated by analyzing statistical data from the experimental velocity pattern (Fig. S8).

In addition, we added some noise to the amplitude of the burst force *f*, as this noise is observed in the experiments (Fig. S9A-D). Hence, we used in our calculations (Eq. 8) force amplitudes (*f*) that are drawn from a Gaussian distribution, centered at *f*_0_, with a constant variance *ψ* = 0.2[*m*][*s*^−2^]. Note that the experimental RF exerts stronger and faster tail bursts as it is chasing a faster VF (Fig. S9E,F). However, for simplicity the simulated RF in our model is only increasing its burst rate in order to chase a faster VF. As we noted before, our main aim was to explore here the implementation of our spin model for directional decision making, so for the modelling of the RF speed, we merely wanted to implement a reasonably realistic model which can be further improved in the future.

The only parameter that we needed to change as a function of the number of VF that the RF is chasing, in order to obtain reasonable agreements with the experimental observations, was found to be *f*_0_: *f*_0_ = 1.1[*m*][*s*^−2^] for 1VF, *f*_0_ = 1.2[*m*][*s*^−2^] in 2VF, and *f*_0_ = 0.95[*m*][*s*^−2^] in 3VF case. It is possible that the RF exerts stronger average tail burst forces when chasing a smaller group of fish since it feels more vulnerable and exposed. Weaker interactions between fish were indeed observed in experiments with larger groups [11]. By reasonable agreement we mean that the simulated and experimental distributions of different quantities that characterize the motion of the RF have the same shape and qualitative features, as well as close average values (see for example the case of 1VF in Fig. 2B-D).

When the RF is following several VF, we need to define how the distance *r* that enters Eq. 9 is calculated. The problem can be framed in terms of which VF the RF pays attention to, at each instant. We define a threshold (*τ*) of fractions of “on” spins (representing the strength of the corresponding neural firing) to determine which targets contribute to the calculation of the distance to targets [14]. On each iteration, the level of neural firing that represents each VF (*n*_*i*_) is monitored to determine if it is above the threshold. If so, the distance to the relevant VF is included in the calculation of *r* as follows

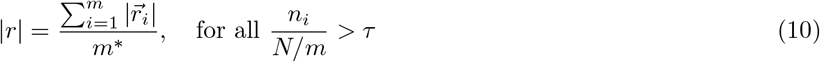

where *m*^*\**^ ≤ *m* is the number of neural groups which fire above the threshold, and 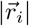 is the distance of the RF to the *i*’th VF (Fig. 1). We used a threshold value of *τ* = 10%, however the results were found not to be very sensitive to the threshold value (Fig. S10).

The model parameters are summarized in table I. Their values are in close agreement with those used in other burst-coast models of this fish motion [22].

**TABLE 1:**
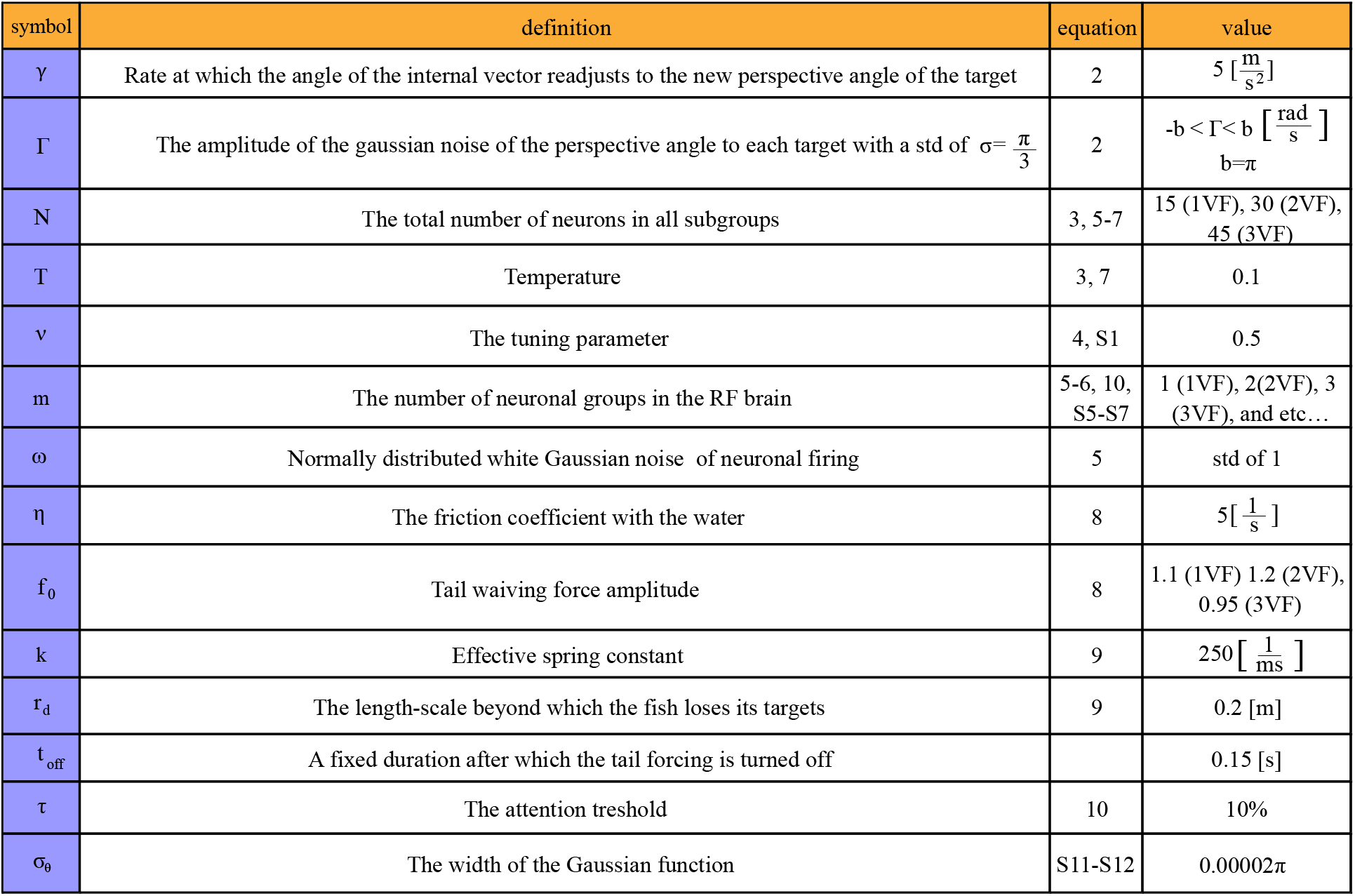
Table of parameters used in our model.

## III. RESULTS

We compare our model with the experiments in which a RF can school with either one, two, or three VF swimming at different average speeds and spatial configurations. We start with calibrating the model for the case of one VF, and then keep all the model parameters fixed when modeling the behavior of RF following two or three VF, except for slightly modulating the mean burst force *f*_0_.

### RF following one VF

We begin with the simplest system of the RF following one VF. A typical time-series of the RF velocity, from the experiment and the simulation, is shown in Fig. 2A. The distributions of the RF speed (*V*_*RF*_, Fig. 2B), maximal speed after each burst (*V*_*peak*_, Fig. 2C), and the time interval between bursts (*t*_*peak*_, Fig. 2D), are all in reasonably good agreement between the experiment and simulations. A typical trajectory of the RF chasing the VF is shown in Fig. 2E,F, in the VF and lab reference frames, respectively (see Supplementary Movie M1).

The accumulated spatial distributions of the RF behind the VF from our simulations are compared to the experimental observations in Fig. 2G-J. The position distributions are shown normalized either over the whole 2D space (Fig. 2G,H), or along individual *x*-axis sections (Fig. 2I,J). We conclude that our model reasonably fits the experiments, for the different mean speeds of the VF. Further comparisons are shown in Fig. S11.

### RF following two VF

In Fig. 3 we plot the accumulated spatial distribution of the RF chasing two VF that are moving at different average speeds, and at different lateral separations (left-right distance, LRD, Fig.1). In Fig. 3A-F we plot the results for a small value of LRD (= 0.06[*m*]), where the experimental data show the RF to be mainly positioned behind, and in-between the 2VF, and no clear bifurcation is observable in the RF distributions (Fig. 3A,C). At larger LRD values (Fig. 3G-L), we clearly see that when the RF is close to the two VF, it follows either one of them, while when it is further away it is found more in the compromise position between them (Fig. 3G,I). Further comparisons are shown in Fig. S12.

**FIG. 3:**
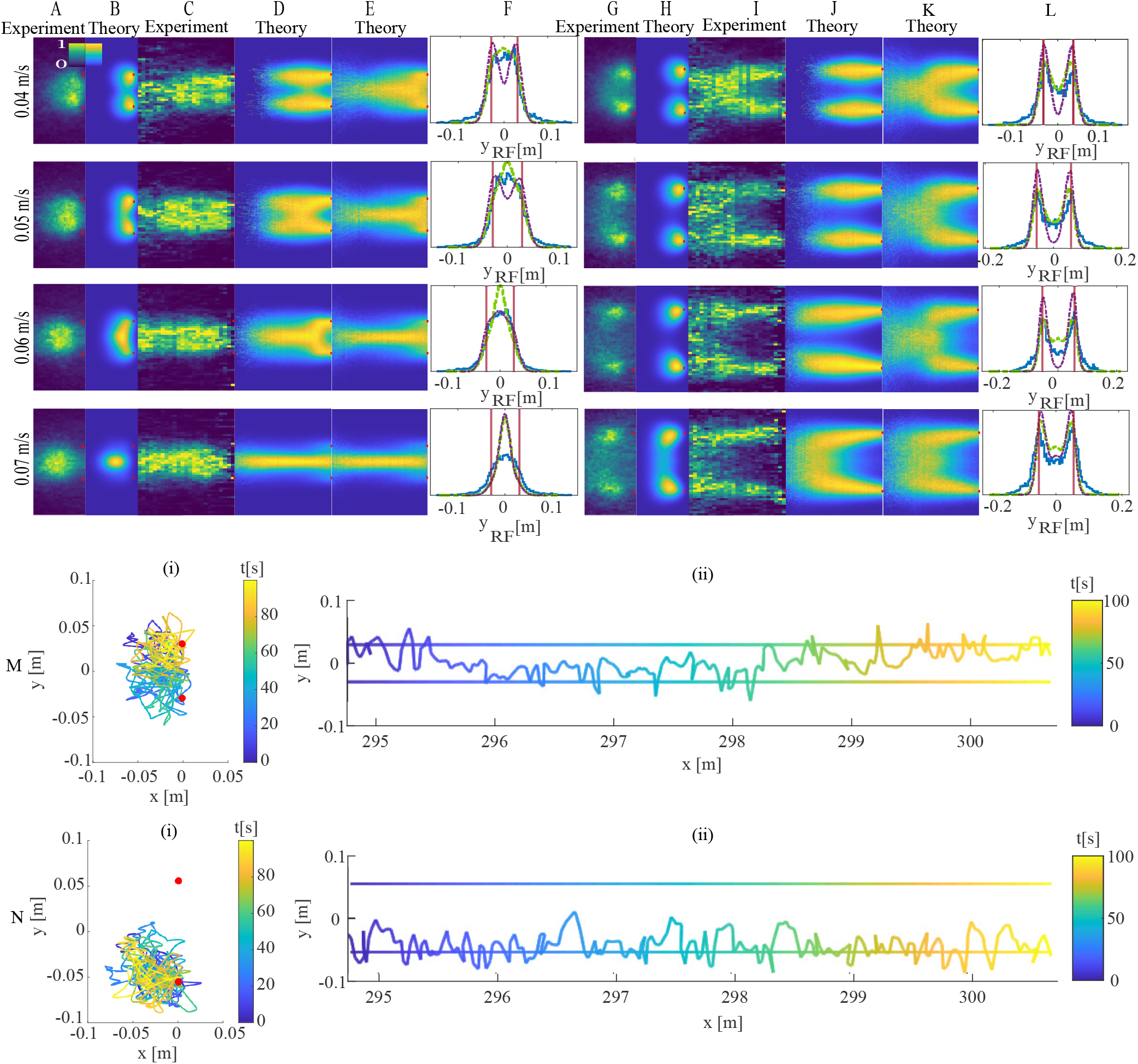
Two VF - spatial distributions of the RF, as heat maps normalized over the whole 2D space (A-B, G-H) or over individual *x*-sections (C-E, I-K). (A,G) and (C, I) shows the experimental RF’s distribution, while (B, H) and (D, E, J, K) show the corresponding simulated distributions (D,J for long simulations, while E,K for short simulations). (F, L) give the distributions of the RF’s *y*-positions and compare the experimental data (blue) with the model (purple for the long simulation and green for the short simulations). The *y*-positions of the VF are denoted by the vertical Maroon lines. (A-F) show results for the compromise regime, in which *LRD* = 0.06*m* for all the VF velocities. (G-L) show the results from the bifurcated regime, in which the LRD between targets is chosen to be 0.08[*m*] for *V*_*V F*_ = 0.04[*m/s*], 0.1[*m*] for *V*_*V F*_ = 0.05[*m/s*], and 0.11[*m*] for *V*_*V F*_ = 0.06[*m/s*], 0.07[*m/s*]). We used the identical parameters as in the 1VF system (Fig. 2), except for: *f*_0_ = 1.2[*m/s*^2^]. We also added the following parameters (for the neuronal tuning and the overlap function): *ν* = 0.5 and *σ*_*θ*_ = 0.00002*π*. The long simulations statistics contain 100 simulations, each ran for 5000[*s*], and the short simulations statistics contains 20, 000 simulations, each ran for 7.5[*s*]. (M) and (N) show typical trajectory of the RF (in a long simulation) in the case of the compromise regime (M), and the bifurcated regime (N), when the VF velocity is 0.06[*m/s*]. M(i) and N(i) show the RF movement relative to the VF, and M(ii) and N(ii) show the movement in the lab frame. For the short simulation in the case of 0.07[*m/s*] the initial condition of *x*_*RF*_ is randomized in the range of 0 − (−0.06)[*m*] behind the VF, while in all other plots it is randomized in the range of 0 − (−0.1)[*m*].

In our model, the motion of the RF between the two VF is termed the “compromise” regime, where the spin dynamics do not break the symmetry. When the symmetry is broken, one group dominates, and the other is inhibited. Such a bifurcation, termed “decision”, arises in our model when the relative angle between the targets, with respect to the RF, is larger than a critical value (of ∼ 90^*o*^ for our choice of model parameters) [14].

Comparing to the simulations, we find that for the small LRD value and low VF speeds (*V*_*V F*_ = 0.04, 0.05, 0.06[*m/s*]), the RF distribution is already bifurcated, i.e. the RF follows behind each of the VF, and not in the compromise position between them (Fig. 3B,D, and Supplementary Movie M2). This behavior is not observed in the experiments, and arises from the fact that the experimental trajectories of the VF and the RF are bounded by the container walls, thereby limiting the experimental data to short trajectories (of ∼ 6 − 8s duration) that are interrupted by U-turns, while the simulated trajectories are long and uninterrupted (around 5, 000[*s*]). We demonstrate this in Fig. 3E,K, by comparing the RF’s spatial distributions obtained from many short simulated trajectories (of 7.5[*s*] duration), that match better the experimental distributions. During short trajectories the spatial distribution of the simulated RF does not reach its steady-state distribution, as obtained from simulating long trajectories.

At the larger LRD values, both the experimental and the simulated distributions exhibit clear bifurcation of the RF dynamics, which follows either one of the two VF (Fig. 3G-L, and Supplementary Movie M3). The projections of the RF’s positions on the *y*-axis (Fig. 3F,L), show that there exists good agreement between the model and the experiments. Similarly, the distributions of the speed, velocity peaks, and intervals between consecutive velocity peaks are in reasonable agreement between the model and the experiments (Fig. S12).

Note that the overlap function did not significantly affect the RF distribution for the simulated two VF system, except at the fastest velocity of the VF (0.07[*m/s*]). This is expected, as the overlap affects the dynamics when the RF is far behind the two VF, so that their relative angle is smaller, which occurs more for the fastest moving VF. At this velocity, without the overlap function, the bifurcation event occurred too close to the targets.

### RF following three VF

The experiments and the simulation results for three VF are displayed in Fig. 4. We find an overall good agreement between the accumulated spatial distributions from the experiments and the corresponding simulations (Fig. 4A-E top two lines). The bifurcations are clearly observed in the model and in the experiments for the larger LRD value (*LRD* = 0.10[*m*]), while for *LRD* = 0.05[*m*] we see the bifurcations clearly only in the simulations but not in the experiments. This is likely due to the larger experimental noise, and to the shorter trajectories in the experiments compared to the long trajectories used in these simulations (as in the 2VF case, see Fig. 3E,K). Further comparisons are shown in Fig. S13, and in Supplementary Movies M4-M5.

The overlap function makes a crucial difference in the dynamics of the RF in the system with three VF, as we demonstrate by plotting the simulated distribution when this effect is not taken into account (Fig. 4A-E bottom line). Without the overlap function, the RF predominantly follows the middle target due to geometrical reasons. Following the middle target appears to be a steady-state solution in the RF decision-making process, in the absence of the overlap normalization, as demonstrated by the sample trajectory (Fig. 4G). In the presence of the overlap function, we observed that the RF chases behind each of the three VF and switches between them (Fig. 4F), and we could also distinguish the bifurcation events on the density maps (Fig. 4A-D, *LRD* = 0.1[*m*]), in agreement with the experimental observations. For small LRD values, the effects of the overlap function are not visible (Fig. S13).

**FIG. 4:**
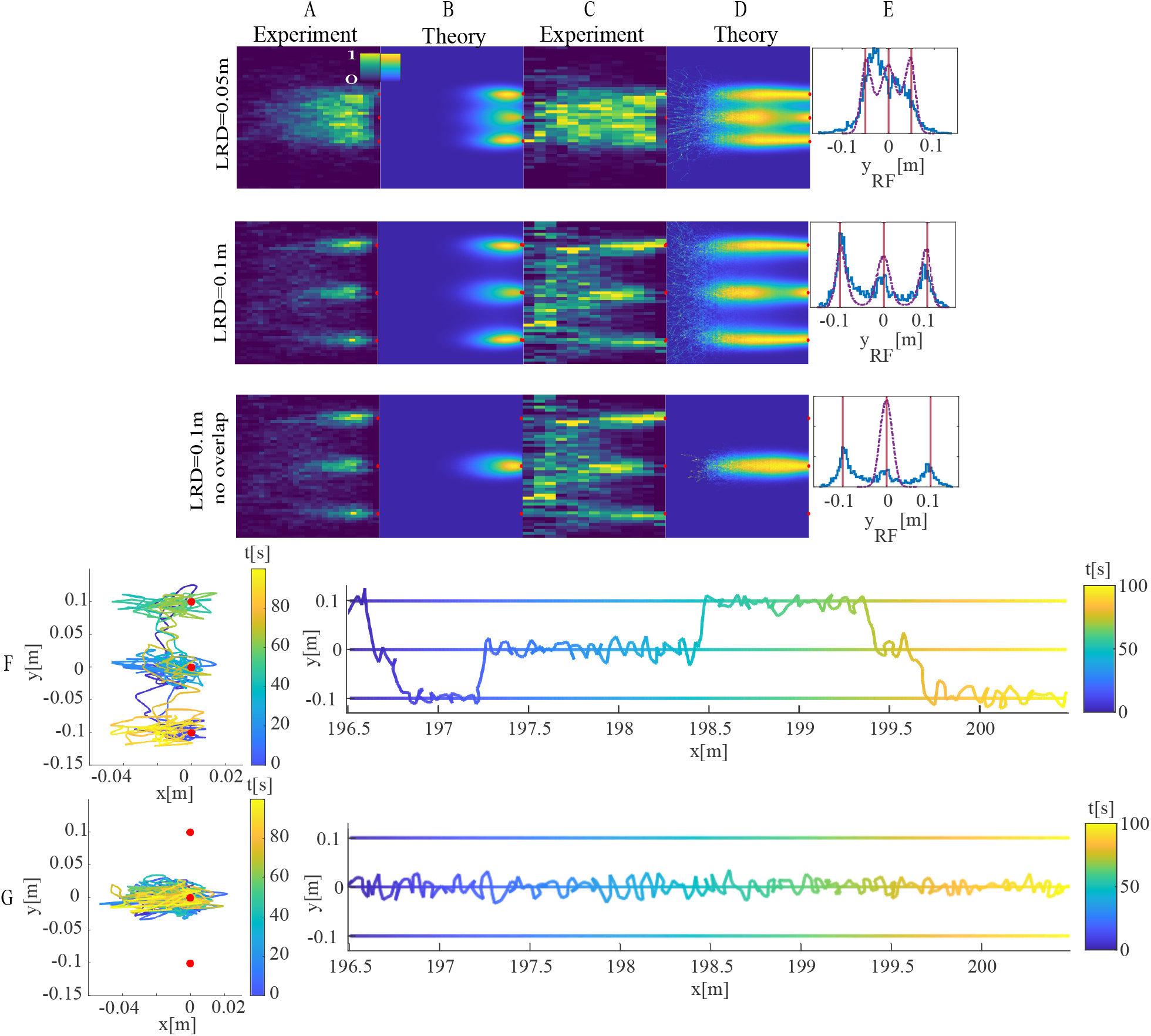
RF following 3VF with an average velocity of *V*_*V F*_ = 0.04[*m/s*]. The value for each line in (A-E) is given on the left, with the third-panel presenting simulations without the overlap function. The heat maps present the accumulated spatial RF distribution, normalized over the whole 2D space (A,B) or over the *x*-axis sections (C,D). (A,C) Experimental data, while (B,D) the simulation results. (E) Distributions of the RF’s projected *y*-positions (experiments in blue and simulations in purple). The *y*-positions of the VF are denoted by the vertical Maroon lines. (F,G) Examples of simulated RF trajectories behind the 3VF when *LRD* = 0.1[*m*] with (F) and without (G) the overlap function. The left panels show the trajectory relative to the 3VF (red dots), and the right panels show the trajectory in the lab frame (trajectories of the 3VF are given by the horizontal lines). Time is given by the color bar. For 3VF cases, we used the same parameters as in 2VF (Fig. 3), except for the burst force amplitude which was changed to *f*_0_ = 0.95[*m/s*^2^]. we ran 100 simulations (in which the RF initial position was random) each for 5000[*s*] (500,000 iteration steps).

### 2VF in shifted configuration

Next, we investigated a configuration of 2VF which are not aligned along the *y*-axis, i.e. along a line that is perpendicular to their direction of motion. One VF was shifted in both the front-back and the left-right directions with respect to the other VF, by a distance of *δ*. The accumulated spatial distributions from both the experiments and simulations are compared in Fig. 5A-D, for different values of the displacement. Overall the model results match the experimental distributions. For example, even in the ‘bifurcation’ regime (*δ* = 0.06[*m*] in Fig. 5A-D), both the experiment and model show that the RF moves between both VF. Typical simulated trajectories demonstrate the motion of the RF between the two VF in Fig. 5E. Further comparisons are shown in Fig. S14, and in Supplementary Movie M6.

**FIG. 5:**
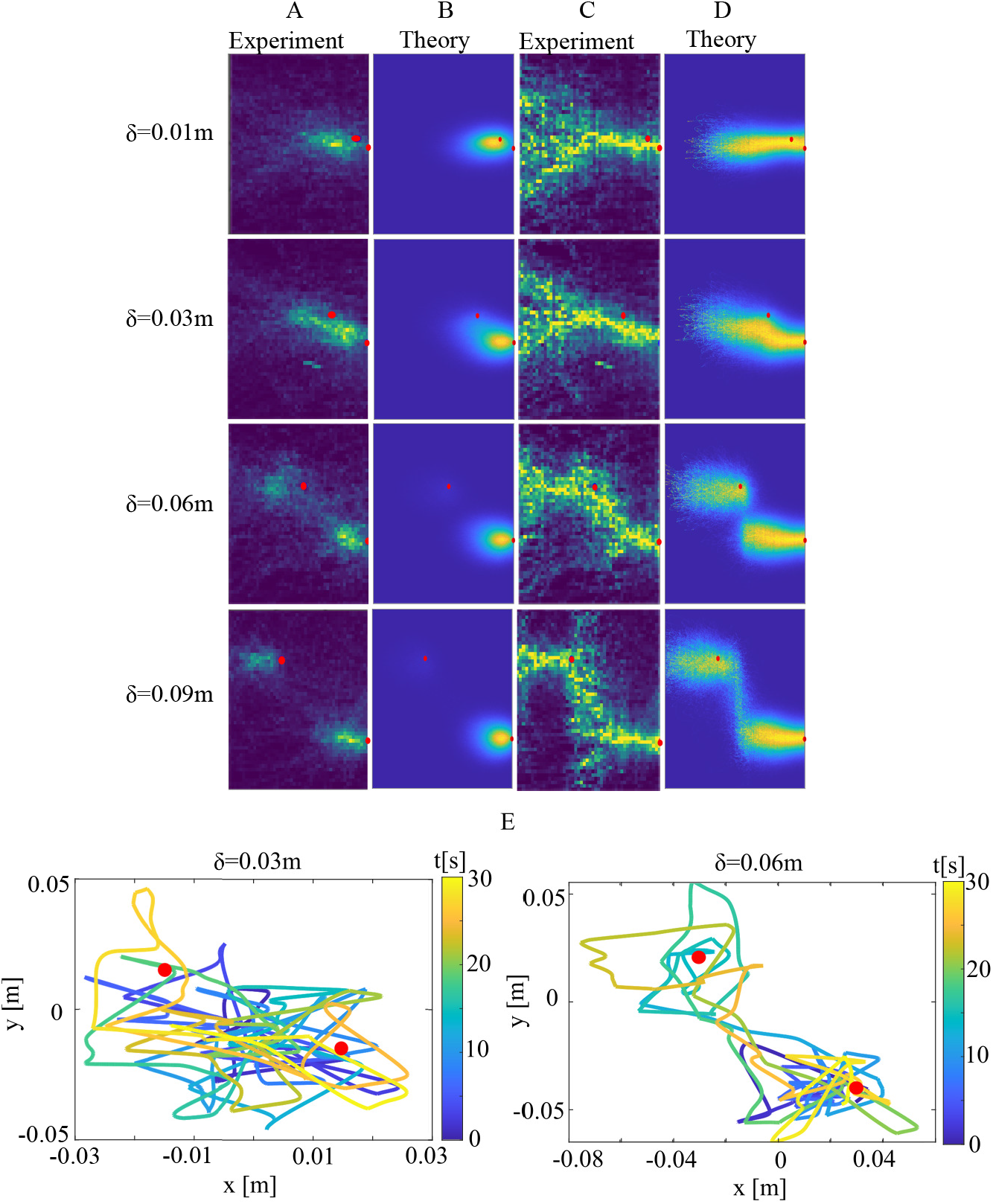
2VF in a shifted geometry: one of the VF is shifted in the front-back and the left-right directions with respect to the other VF, by a distance of *δ*. Both of the VF swim with an average velocity of *V*_*V F*_ = 0.04[*m/s*]. The accumulated spatial distribution of the RF behind the 2VF, (A,B) normalized over the whole 2D space, or (C,D) normalized over *x*-axis sections. (A) and (C) show the results from experimental data, while (B) and (D) show results from the simulations. The VF are denoted by the red circles. The model parameters are the same as those used in Fig. 3. (E) Examples of RF trajectories relative to the 2VF (red dots), for the cases of *δ* = 0.03[*m*], and *δ* = 0.06[*m*]. In both plots, the color of the line represents the time, as indicated by the colorbar, indicating that the RF can go back to follow the trailing VF.

This agreement demonstrates that our model is not limited to describing a fish responding to a single line of leaders, but can also account for the fish motion within a moving group with complex geometry. Note that the one-dimensional version of the model, used in [14], cannot explain the trajectories in this configuration.

## IV. DISCUSSION

Our research provides a novel approach to modeling the motion of fish when responding socially to other fish, by including an explicit model for the directional decision-making processes of the individual fish. This is implemented using a spin-based model [14], which gives rise to spontaneous symmetry breaking, describing the cognitive process of choosing which of the targets to follow. Previous models of animals following others moving in a group usually rely on alignment interactions or vectorial averaging, which do not contain a mechanism for choice. While the parameters used in our model simulations are specific to the zebrafish (*Danio rerio*) that were examined in the experiments, the proposed directional decision-making model may apply more generally to other animals that move in groups.

The Ising spin model that we used for the directional decision-making, is shown to provide an appropriate framework for representing the fish’s directional decisions while it is chasing moving targets. While previously applied under the constraints of projecting it to a 1D motion [14], in this work we have extended it to describe the free movement of the fish in 2D when following other moving conspecifics. The model simulations recovered the main dynamical features of the experimentally observed RF motion. Most importantly, the model captures the observed bifurcation transition exhibited by real fish, as a function of the separation between the VF: when the VF are close to each other, the RF follows them along an averaged (compromise) position, while when the VF are far apart, the RF pursues each of them individually. Moreover, we used identical parameters for VF of different speeds and geometries, indicating that the model is rather general and robust.

Our model may form a first step towards implementing individual directional decision-making in models of animal collective motion. As shown in Fig. S15, we can explore the dynamics of many RF interacting using our model, following a single leader, or in its absence. These results indicate that the model can describe the formation of cohesive shoal (“swarm”) behavior, without any spontaneous alignment of the fish along a particular direction [26] (see Supplementary Movie M10). This suggests that for the maintenance of collective persistent motion along a particular direction (“schooling phase” [7, 13]), the spontaneous emergence of leaders is needed [27–29], such as leadership by indifference (whereby less socially-responsive individuals spontaneously become leaders [27]), which is not included in the current model. Note that our model ignores the finite size of the RF, which means that we do not describe the short-range maneuvers that are necessary for collision avoidance. Similarly, in a real fish school some individuals may have more influence than others [4, 30]. In addition, our model explicitly involves many-body interactions, that are not simply the superposition of pairwise interactions. Significant interactions beyond pairwise have indeed been measured inside fish schools [11].

The model can be elaborated and developed to include more detailed aspects of the interactions between the fish [3, 4, 8, 13, 30, 31], while the explicit directional decision-making model may be incorporated into existing models of collective animal motion [32, 33]. It remains for future exploration how this individual decision-making process can give rise to the emergence of polarized fish schools and (possibly transient) leaders.

## V. ACKNOWLEDGMENTS

We thank Pawel Romanczuk for useful comments. N.S.G. is the incumbent of the Lee and William Abramowitz Professorial Chair of Biophysics. This research is made possible in part by the historic generosity of the Harold Perlman Family. D.G. and I.D.C. acknowledge support from the Office of Naval Research Grant N0001419-1-2556, Germany’s Excellence Strategy-EXC 2117–422037984 (to I.D.C.) and the Max Planck Society, as well as the European Union’s Horizon 2020 research and innovation programme under the Marie Skłodowska-Curie grant agreement (to I.D.C.; #860949).

