## Supplementary material for "A simple cognitive model explains movement decisions during schooling in zebrafish"


### S1. TUNING PARAMETER OF THE RELATIVE ANGLES

We investigated a modified form of the neuronal interactions based on the angle between its targets. The interaction between the spins of different sub-groups uses a modified angle according to:

$$\theta_{ij} \to \theta^*_{ij} = \pi \left( \frac{|\theta_{ij}|}{\pi} \right)^{\nu} \tag{S1}$$

where $\theta_{ij}$ is the original measured conflict angle (as shown in Fig.1), $\theta^*_{ij}$ is the new modified angle and the tuning parameter $\nu$ describes the shape of the interaction between neural circuits, corresponding to the wiring or filtering activity of the brain's neurons (Fig.S1). Hence the interaction between the spins was modified to:

$$J_{ij} = cos(\theta^*_{ij}) \tag{S2}$$

From previous experiments, a suitable tuning parameter to represent animal movement was found to be $\nu \approx 0.5$ [1], therefore we will use this value in our simulations. In Fig.S1 we plot the affect of $\nu$ on the conflict angle in which excitation ($J_{ij} > 0$) changes to inhibition ($J_{ij} < 0$).

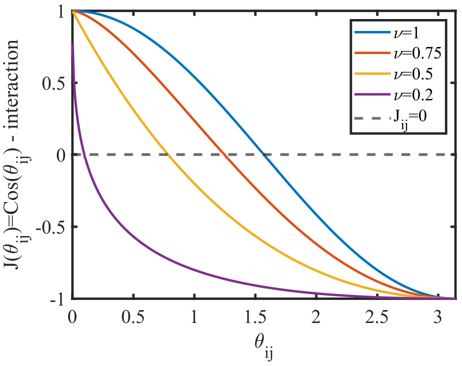


FIG. S1: The tuning parameter affect the conflict angle between targets ($\theta_{ij}$) in which we get inhibition ($J_{ij} < 0$) or excitation ($J_{ij} > 0$). As the tuning parameter is smaller, the conflict angle needed to inhibition is smaller. We used $\nu = 0.5$ (in orange) in our fish model.

### S2. CALCULATING THE NEURONAL NOISE TERM IN THE CONTINUUM MODEL

To calculate the neural noise which we added to the continuum approach, for each subgroup of neurons, we first defined $r_-$ and $r_+$ in each group to be:


*

$$r_+ = k_{i,on}(\frac{N}{m} - n_i)$$
$$r_- = k_{i,off} n_i \tag{S3}$$

where, $m$ is the number of neural groups (and also the number of VF), and $n_i$ is the amount of "on" neurons in group $i$. By substituting the Glauber rates as defined in Eq.7 in Eq.S3, we get:

$$r_+ = \frac{\frac{N}{m} - n_i}{1+\exp\left(-\frac{n_i + \sum_{j\neq i}^{m} n_j \cos\theta_{ij}}{T}\right)}$$
$$r_- = \frac{n_i}{1+\exp\left(\frac{n_i + \sum_{j\neq i}^{m} n_j \cos\theta_{ij}}{T}\right)} \tag{S4}$$

We will focus on one event change at most in $\delta n_i$:

$$\delta n_i = \begin{cases} +\frac{1}{\frac{1}{m}N} & P_+ = r_+\delta t \\ -\frac{1}{\frac{1}{m}N} & P_- = r_-\delta t \end{cases} \tag{S5}$$

here $P_+$ and $P_-$ are the probabilities of the neurons to turn "on" or "off, and the variance in neural firing is:

$$\delta n_i^2 = \frac{P^+ + P^-}{\left(\frac{N}{m}\right)^2} = \frac{r_+\delta t + r_-\delta t}{(\frac{N}{m})^2} \tag{S6}$$

Next, we calculate the neural noise amplitude, $B(n_i)$:

$$B(n_i) = \lim_{\delta t \to 0} \frac{< \delta n_i^2 >}{\delta t} \tag{S7}$$

and by substituting $\delta n_i^2$ inside the previous equation, we get:

$$B(n_i) = \frac{r_+ + r_-}{\left(\frac{N}{m}\right)^2} \tag{S8}$$

We will substitute Eq.S4 in Eq.S8 and this will be calculated each time step to find the neural noise amplitude which depend on $N$, $T$, $n_i$, $n_j$, and $\theta_{ij}$:

$$B(n_i) = \frac{k_{i,on}(\frac{N}{m} - n_i) + k_{i,off} n_i}{\left(\frac{N}{m}\right)^2} \tag{S9}$$

We used Eq.S9 as the amplitude of the neural noise as follows:

$$\frac{dn_i}{dt} = \left(\frac{N}{m} - n_i\right) k_{i,on} - n_i k_{i,off} + \omega\sqrt{B(n_i)} \tag{S10}$$

$B(n_i)$ is the noise amplitude, and $\omega$ is a normally distributed white Gaussian noise (with a standard deviation of 1 in our model).

### S3. THE OVERLAP FUNCTION

The main objective for the overlap function is to represent a situation in which several VF are within very close viewing angles with respect to the RF eyes. In such cases, we want the RF to "see" them as "one" target, so the neural groups will not fire fully to both VF, but will overlap with each other. This will decrease the total neurons that can be turned "on" in each group. In this manner, the fish will not tend to follow the overlapping targets more than isolated VF, since it sees them as approximately one fish. Figure S2 illustrates such a case in which the overlapping function could affect the neural firing of groups number 1 and 2 but does not change the dynamics in group number 3. The RF sees both $VF_1$ and $VF_2$ at very similar angles, and therefore their relative neural groups will overlap and will be reduced.

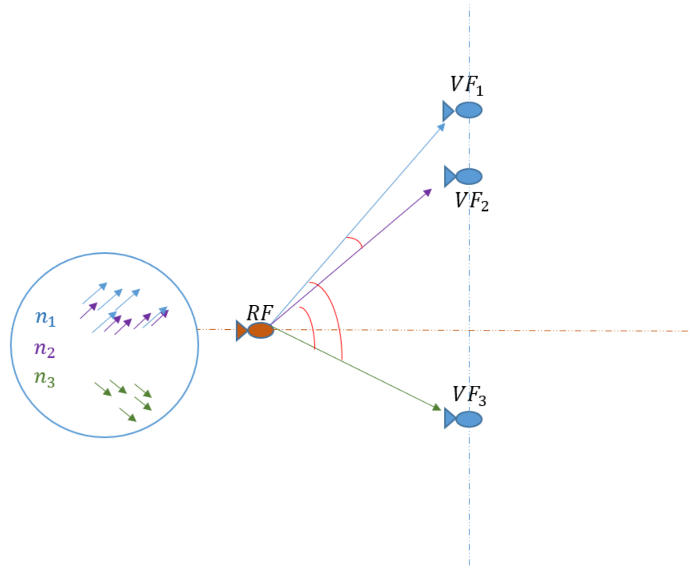


FIG. S2: The Overlap function in cases of overlapping targets - The RF observes $VF_1$ and $VF_2$ at very similar angles, yet it sees $VF_3$ at a distinct angle. Therefore, the overlap function for the RF has been employed to avoid bias to the direction of the $VF_1$ and $VF_2$. This decreases the total amount of neural firing in $n_1$ and $n_2$ depending on how the related angles overlap (how close is $\theta_1$ to $\theta_2$). The $n_3$ group may also be affected, but to a much lesser degree.

Each set of spins ($n_i$) is aimed at angle $\theta_i$ which points towards target $i$, but for simplicity we assume it has a Gaussian form around this angle:

$$f_i(\theta_i, \theta) = A \exp^{-\frac{(\theta-\theta_i)^2}{\sigma_\theta}} \tag{S11}$$

Where $\sigma_\theta$ is the width of the Gaussian function, and $A$ is a normalization factor such that: $\int_{\theta_i-b}^{\theta_i+b} f_i(\theta_i, \theta) d\theta = 1$ (the limits of the integral $b$, will be given next). To demonstrate the overlap functions, we present Fig.(S3).

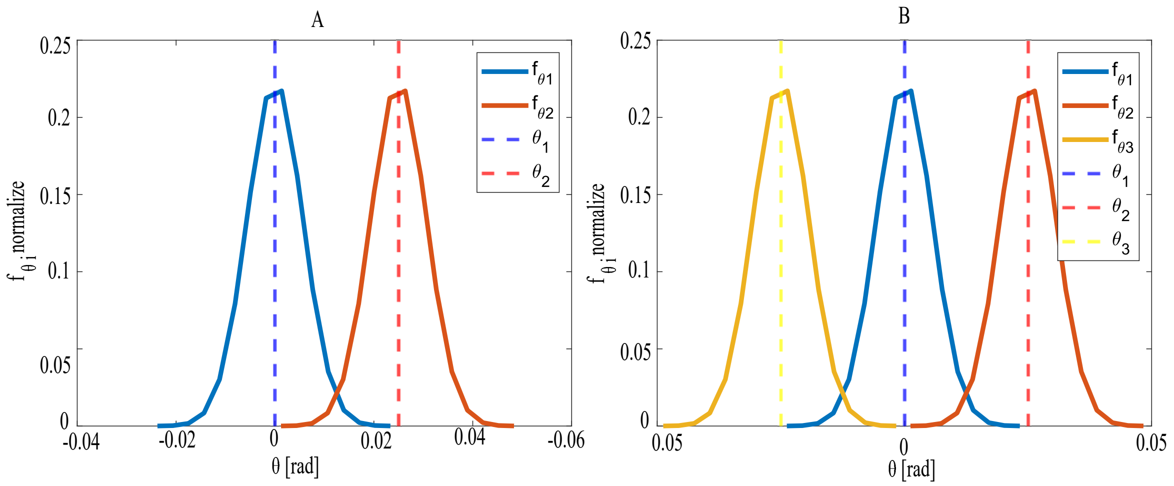


FIG. S3: The Gaussian function of Eq.S11 representing the neuronal spread of the neurons that point to each target. The right plot (A) shows two targets configuration (2VF), and the left plot (B) shows three targets configuration (3VF). The 3VF is symmetrical around the angle $\theta_1$ directed at the central target. We chose a value of $\sigma_\theta = 0.00002\pi$ for both 2VF and 3VF. The values of the normalization factor in these examples are: in 2VF - overlap function of groups 1 and 2 get 0.9805. In 3VF-overlap function of groups 1 and 2 is 0.9805 and the overlap function of group 3 equals 0.9610.

The normalization of the neuronal strength due to the overlap function is calculated in the following manner:

$$n_i = n_{i.0} \int_{\theta_i - b}^{\theta_i + b} \frac{f_i(\theta_i, \theta)}{\sum_j f_j(\theta_j, \theta)} f_i(\theta_i, \theta) d\theta \tag{S12}$$

Here $b = 3\sqrt{\sigma_\theta}$, $n_{i,0}$ is the original neural firing before including the overlapping, $f_i(\theta_i, \theta)$ is from Eq.S11 and $f_j(\theta_j, \theta)$ is the same just for other VF, in the direction of $\theta_j$. The sum in the denominator goes over all the subgroups (to each VF), including $j = i$. At every angle $\theta_i$ the fraction of the spins of group $i$ that are counted is not $f_i(\theta_i, \theta)d\theta$, but decreased due to the overlap with the other Gaussian functions around the $\theta_j$, as given by the normalizing factor $\frac{f_i(\theta_i,\theta)}{\sum_j f_j(\theta_j,\theta)}$.

In 2VF, when the VF's angles overlap ($\theta_1 - \theta_2 \rightarrow 0$), the function goes to $\frac{1}{2}$, and we do not double-count the spins that point at the same target. When the targets do not overlap, each recieves its full share of $n_{i,0}$ spins. In 3VF case, there can be symmetry around the middle target ($\theta_1$). When the three targets overlap, the overlap function of each of them goes to 1/3 so that we do not triple-count the spins that point at the same target. The central target is affected by a stronger overlap since it is affected by both edge targets.

We kept the parameter $\sigma_\theta$ equal to $0.00002\pi$ on our simulations. We confirmed that the overlap contributes to our model in Fig.S13. Before appending the overlap function, the RF only chases behind the middle target of the 3VF (Fig.S13). However, when the overlap is included, the RF chases behind all three VF, which better fits the experiment. The reason is that the overlap causes the middle VF to have lower amplitude due to overlapping with the nearby VF from both sides. Subsequently the neural firing of the group directed at the middle target decreases.

### S4. EFFECTIVE INTERACTION BETWEEN THE RF AND VF, EXPERIMENTS AND MODEL

The relative acceleration of a fish following other fish, or another VF, was measured in experiments (Fig.S4A-D). A similar analysis of our simulations is shown in Fig.S4E,F.

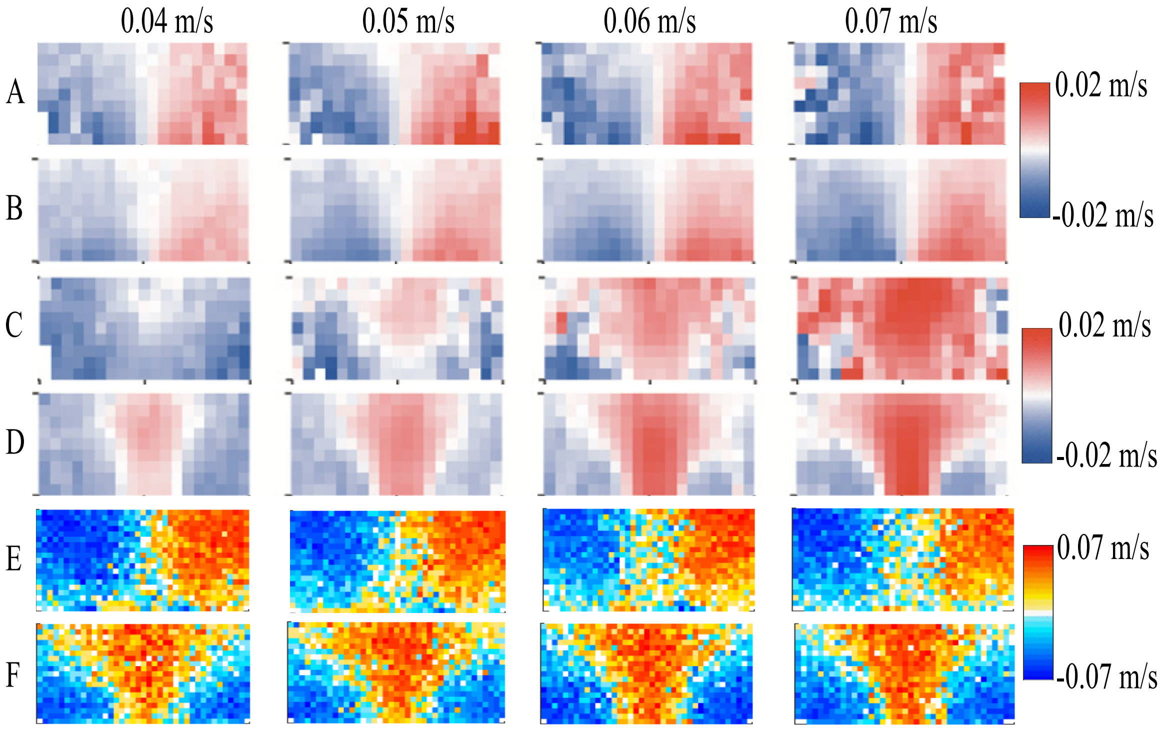


FIG. S4: The effective interactions between fish, as manifested by their acceleration as function of their position in relation to the fish they are chasing. The chasing RF is at the origin $(0, 0)$ and the target is at different positions around it. The $x$-axis is the front back distance between the RF and its target, $y$-axis is the relative left/right displacement. The turning speed and forward speed components are shown by the colormap. (A) Experimental data showing turning components of the RF's speed when chasing another RF, or a VF (B). (C) Experimental data showing forward components of the RF's speed when chasing another RF, or a VF (D). (E-F) The simulated turning (E) and forward speed (F), using the same parameters as in Fig.2.

### S5. SPRING-LIKE MODEL FOR THE RF SPEED

Before applying the tail burst events in our model, we tried a simpler approach, where the velocity amplitude is directly calculated in a continuous manner from the distance of the RF to its targets (depending on the attention of the RF). The RF velocity amplitude is calculated in the following way:

$$V_0(|\vec{r}|) = k|\vec{r}| \exp\left(\frac{-|\vec{r}|^2}{2r_d^2}\right) \tag{S13}$$

where $\vec{r} = \vec{r}_{rf} - \vec{r}_{vf}$ is the absolute distance of the RF from the VF, $V_0$ is the speed value, $r_d$ is a constant that gives the range beyond which the interactions between the fish decay, and $k$ is the spring constant. We added a finite time scale that adjusts the speed of the RF in order for it to reach closer to the VF as in experiments. Together with a noise in the velocity of the RF, such that the velocity distribution is wider and fit better. Both were added by applying the following equation:

$$\frac{dV}{dt} = -\beta(V - V_0) + \xi \tag{S14}$$

where $\beta$ is the frequency to adjust the velocity, and $\xi$ is a white Gaussian noise term.

We apply this spring like method on the the two dimensional system for one, two and three targets (Fig.S5) This model resulted in a reasonably good agreement with experimental heat maps (Fig.S5A-E). However, the RF's velocity dynamics did not match the velocity pattern of the RF in experiments (Fig.S5F). Therefore, we focused in the main text on a model of realistic burst-and-coast behavior of the RF. The spring-like model did not give realistic spatial distributions when the 2VF where in a shifted geometry (Fig.5).

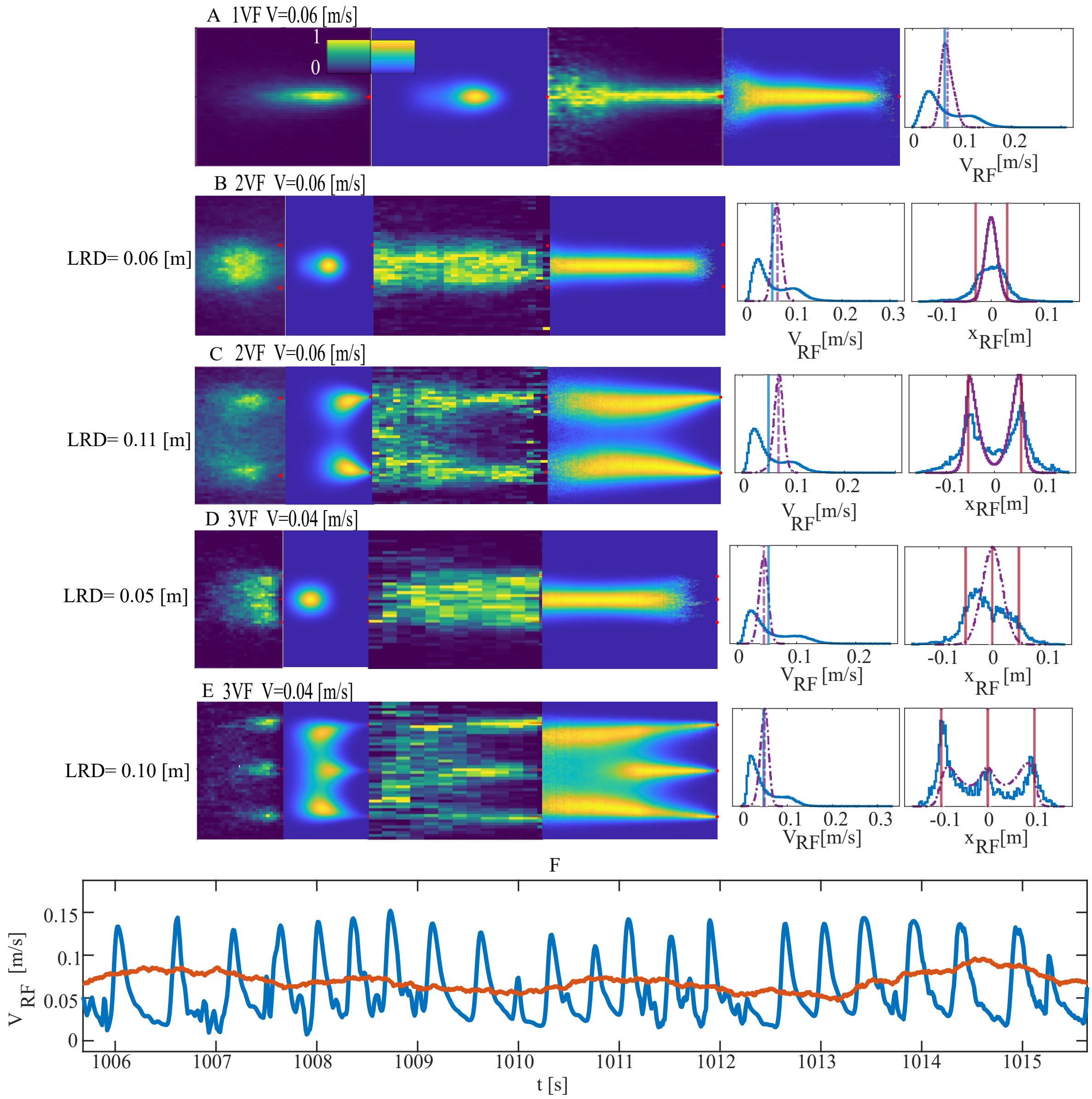


FIG. S5: Spring Like interactions model - Heat maps of the spatial distribution of the RF chasing (A) 1VF, (B,C) 2VF and (D,E) 3VF. The right panels give the speed, and projected $y$-position distributions. The blue/purple line show the experimental/simulated distribution. The bottom plot show the RF dynamics in experiment (blue), and the red line the RF velocity from the spring-like simulation. For good statistics we conducted for each heat map and histogram 500 simulations, with random initial position of the RF, each for $5,000[s]$ which equal to 500,000 iteration steps.

### S6. BURST EVENTS

In order to simulate the tail bursts we used a Gillespie method to determine whether a tail burst just occurred or not, depending on the rate of the tail to exert a burst (Eq.9). The Gillespie simulation determines in each iteration whether a tail burst just happen ($F = 1$) or stopped ($F = 0$). The first step is to determine the time of the next event, a number $r$ is drawn from a uniform distribution and the next time step is calculated by:

$$dt = \frac{1}{k_{s,on}} log\left(\frac{1}{r}\right) \tag{S15}$$

where $k_{s,on}$ the rate to burst, as given in Eq.9. We defined the time step between iterations to be $\Delta t = 0.01[s]$, so that during each iteration we check whether $dt < \Delta t$ and also if $V_{RF} < V_{threshold}$ (see section S8 below). If both of these conditions are satisfied, a tail burst occurs and $F = 1$, which is then inserted in Eq.8, and affects the RF's velocity.

### S7. THE BURST RATE VS THE DISTANCE TO TARGET

In order to examine if the experimental tail bursts are more frequent as the distance to the target fish is larger, as assumed in our model (Eq.9), we collected all the bursts events (the minimums of the $V_{RF}$) and binned them as function of the distances ($|\vec{r}|$) in which they occur (Fig.S6, $P(burst)$). The rate to give a burst is calculated as follows: we first computed the probability to be at each location (by excluding all the locations just after the bursts and until the maximum of the velocity), and then divided the probability to burst at each distance ($P(burst)$) by the probability to be at each distance ($P(r)$), and get the rate of the tail to burst at different distances:

$$f_{burst}[\frac{1}{sec}] = \frac{P(burst)}{P(r)\Delta t}$$

where $\Delta t = 0.01[s]$ is the time step and is the same in both the experiments and the simulations. To check whether the frequency is indeed higher at greater distances, we plot the rate ($f_{burst}$) for different distances, $|\vec{r}|$ (Fig.S6), and we concluded that the frequency to exert a burst is higher for greater distances. However, the statistics from the experiments are pretty low, due to the short duration of the experimental trajectories. Also, the probability to be at edge positions, very far from the target (above $0.07[m]$) or very close to the target (below $0.02[m]$), is low, and still, we may get a burst event in those positions, resulting in large fluctuations in the calculated burst rate (dividing by a small number in Eq.S7). For the fastest VF, the linear relation is clear in the experimental data, while it is not very clear for the slower moving VF.

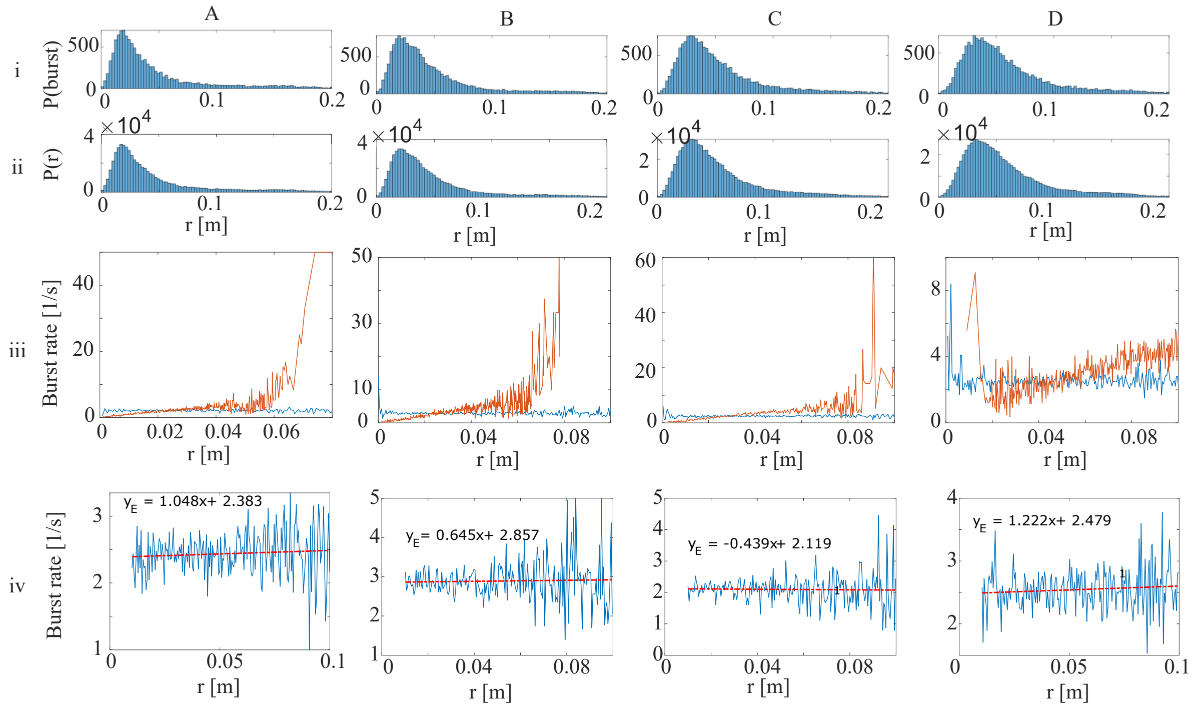


FIG. S6: Rate to give a burst as function of the distance to the VF. (A) for $V_{VF} = 0.04[m/s]$, (B) for $V_{VF} = 0.05[m/s]$, (C) for $V_{VF} = 0.06[m/s]$, and (D) for $V_{VF} = 0.07[m/s]$. First row of plots show the probability to be at each distance, $P(r)$, and the probability to burst at each distance, $P(burst)$ (for the experimental data). For calculating the $P(burst)$ we collect distances where we have a burst, then bin the distances to 100 bins, and for each of them calculate how many bursts occurred. Then we calculate $P(r)$ - the probability that the RF spend time in each of our bins, and we excluded from it the positions during the bursts (the time between burst and next peak of speed). The middle row shows the calculated burst rate, using Eq.S7, in experiments (blue) and the model simulations (orange). The last row of plots present a linear fit to the experimental rate to burst vs the distance to the target.

### S8. VELOCITY THRESHOLD

In order for the simulated tail bursts to occur in a realistic manner, we found it crucial to define the velocity threshold. Only when the RF's velocity is below the value of the threshold ($0.04[m/s]$), a tail burst could occur (meaning we randomize a number - the tail burst event, as described in the previous section). An example of the tail waiving model without such a threshold is presented in Fig.S7, in which it is clear that the tail bursts could happen too quickly from the previous peak without this threshold (see the purple line).

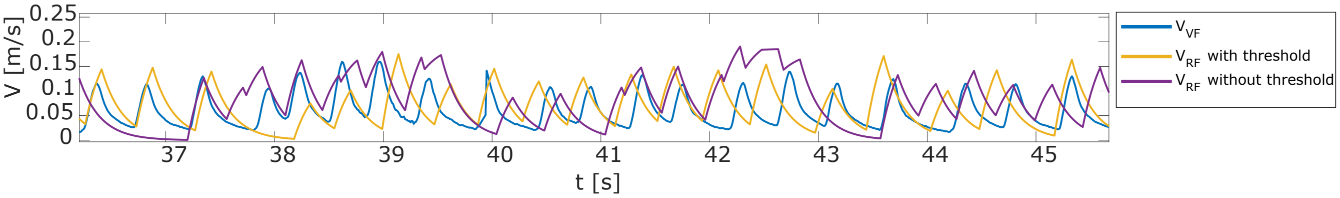


FIG. S7: The RF velocity: the simulated RF's velocity dynamics without the velocity threshold is shown in purple, while the RF's velocity with the threshold ($0.04[m/s]$) is in yellow. The experimental VF velocity is in blue. We used here the same parameters as in Fig.2 (average $V_{VF} = 0.06[m/s]$).

We demonstrated why we chose a velocity threshold of $0.04[m/s]$ in Fig.S8, where are shown histograms of the experimental RF speed values at the burst events. The values of minima of the speeds (where the burst events occur) are mostly below our chosen threshold. When we tried a higher value the RF velocity dynamics didn't match the experiments.

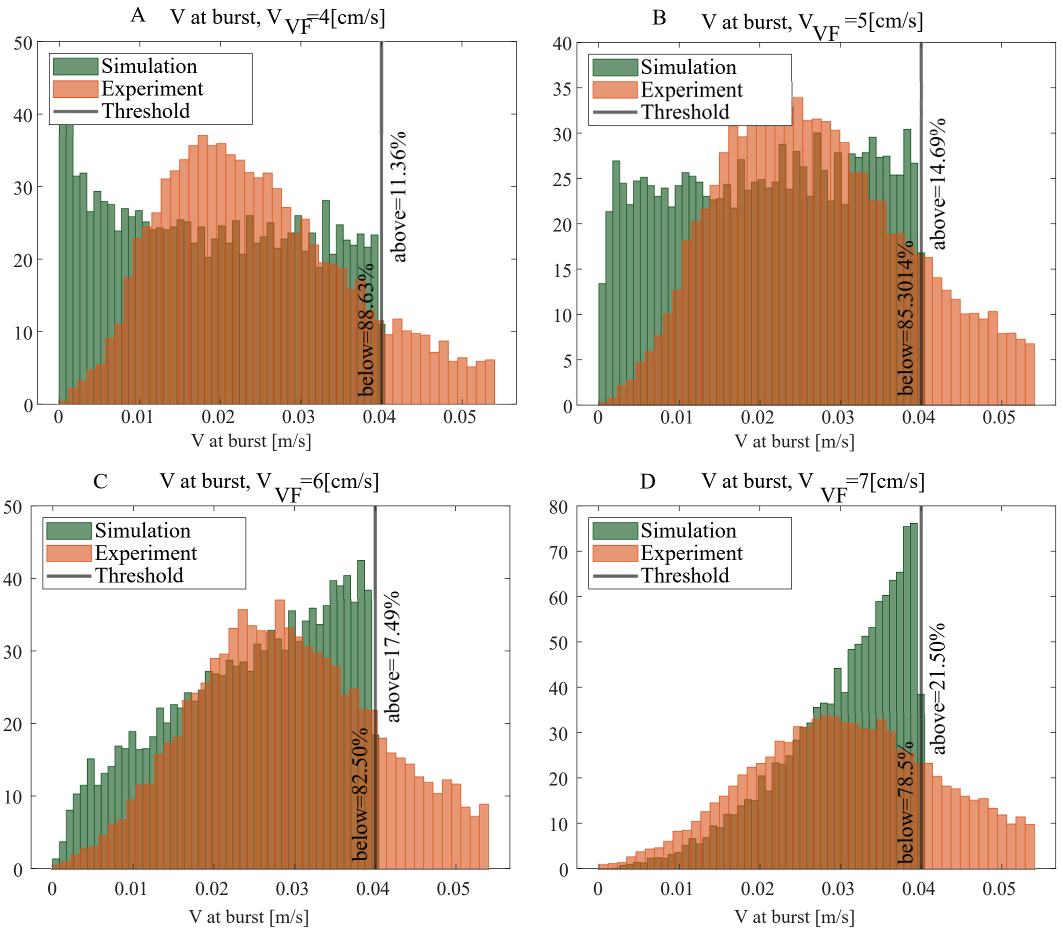


FIG. S8: The distributions of the velocity at the onset of bursts for different VF velocities. The green distributions are for simulations and the orange distributions for the experiments. The black line is the velocity threshold we chose ($0.04[m/s]$), below which most of the experimental bursts occur.

### S9. DISTRIBUTION OF THE AMPLITUDE OF THE VELOCITY BURSTS FROM THE EXPERIMENTAL RF DYNAMICS

We extracted the difference in the experimental RF velocity between the time of burst initiation and its maximum.

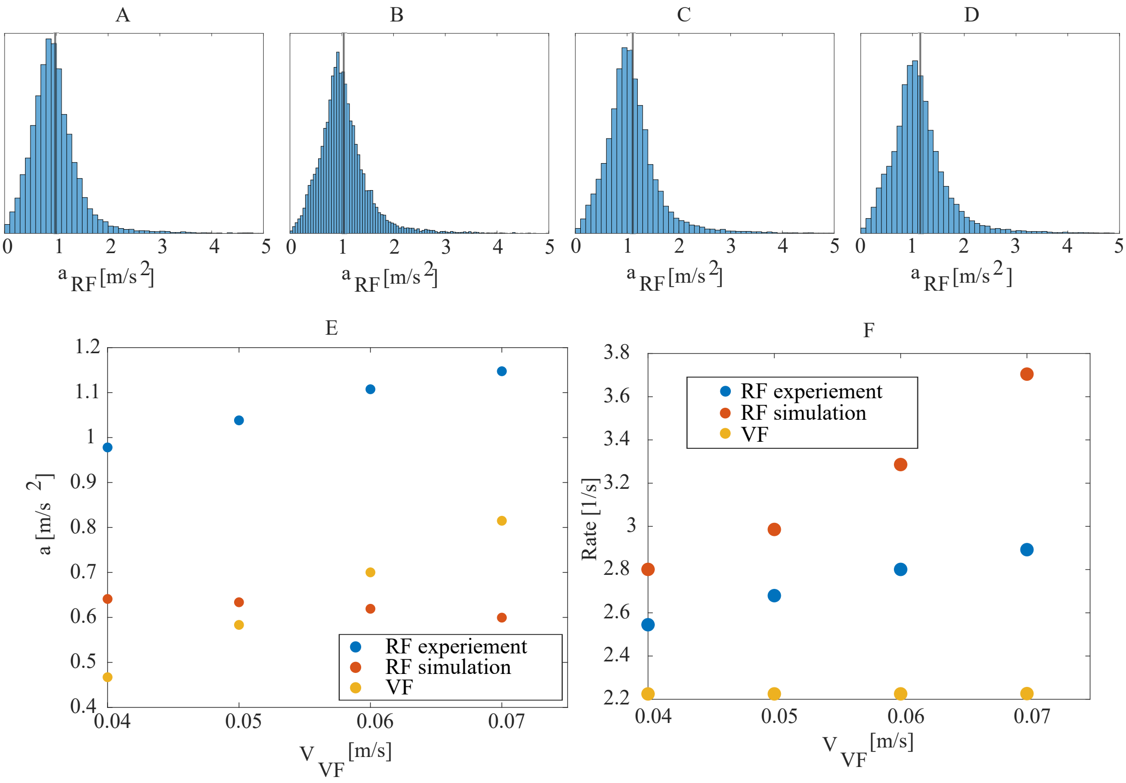


FIG. S9: Distributions of the RF's acceleration (burst amplitude divided by the time of each burst) from the experimental data, following 1VF: (A) $V_{VF} = 0.04[m/s]$, (B) $V_{VF} = 0.05[m/s]$, (C) $V_{VF} = 0.06[m/s]$, and (D) $V_{VF} = 0.07[m/s]$. (E) The average acceleration amplitude for the experimental RF (blue), simulated RF (red), and the VF (yellow). (F) The average rate of the tail bursts for the RF in experiments (blue), the simulated RF (red), and the VF (yellow). This rate is calculated as the average time duration between a burst event and the timing of the prior peak velocity.

### S10. ATTENTION THRESHOLD OF THE RF

By expanding the model from 1VF to several VF, we defined a threshold ($\tau$) of neural firing to determine which targets contribute to the calculation of the distance to targets, Eq.9. On each iteration, the neural firing ($n_i$) is monitored to determine if it is above the threshold. If so, the distance to the relevant VF is included in the calculation of $|\vec{r}|$ as follows:

$$|r| = \frac{\sum_{i=1}^{m} |\vec{r}_i|}{m}, \quad \text{for all } \frac{n_i}{N/k} > \tau \tag{S16}$$

where $m$ equals the number of neural groups which fire above the threshold. The $N$ is the total number of neurons in all the groups, $k$ is the number of neural groups, and $|\vec{r}_i|$ is the calculated distance of the RF to target $i$.

In the case of the 2VF and 3VF, we included: $\tau = 10\%$. As shown in Fig.S10 the RF's $y$-position behind the targets are not very sensitive to the threshold value.

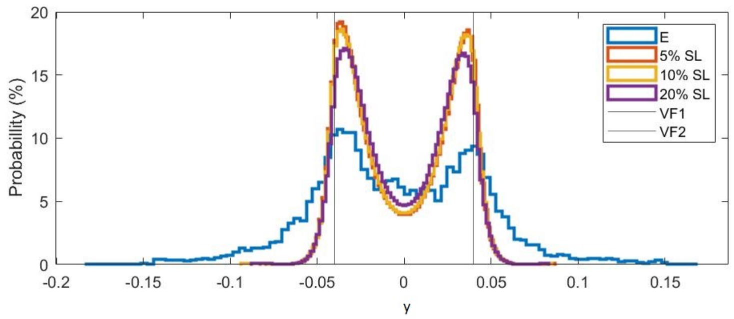


FIG. S10: Examining the effect of different $\tau$ parameters on the probability distrubutions of $y_{RF}[m]$, the simulations ran for $V_{VF} = 0.04[m/s]$. Using the following parameters: $\beta = 1[1/s]$, $\gamma = 5[1/s]$, $\sigma = \pi[rad/s]$, $\xi = 0.01[m/s^2]$, $b = \pi[rad/s]$, $\nu = 0.5$, $r_d = 0.2[m]$, $\sigma_\theta = 0.00002\pi$, and $k = 0.95[1/s]$.

### S11. RF FOLLOWING 1VF

We present here extended data for the case of the RF following 1VF (complimentary to Fig.2). For each velocity we compare the distribution of the distance between the RF and the VF along the $x$-axis (direction of motion of the VF, Fig.S11E), which emphasizes that the RF lags further behind the VF as the VF speed increases. In our model this trend arises, as the tail-bursts occur more rapidly as the distance to the VF increases (Eq.9), enabling the RF to chase faster VF at larger lag distances. At the fastest VF speed the model gives a lag distance that is significantly larger than the experiment. However, all other measures of the RF dynamics, as shown in Fig.S11F-H, are in reasonable agreement across all VF speeds. Although a more precise fit could be obtained by adjusting the mean burst force $f_0$ parameter to each $V_{VF}$, we prefer to keep all the model parameters as constant, such that the model is as general as possible.

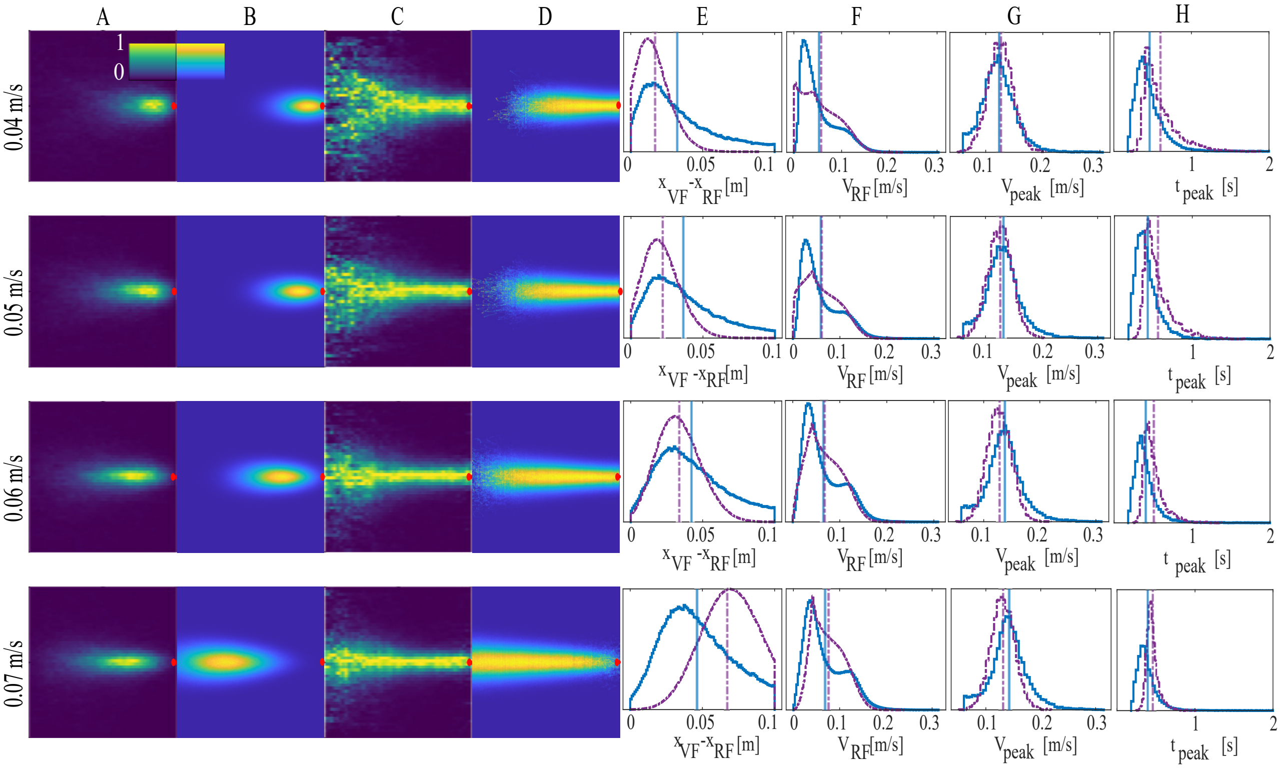


FIG. S11: RF chasing 1VF. (A-D) Accumulated distribution of the RF behind the VF, for different mean $V_{VF}$ (each row), in the VF frame of reference (the VF is at the origin, red dot). For each velocity we normalized the heat maps over the whole 2D space (A and B), or over individual lines of constant $x$-sections (C and D). (A) and (C) Displays the experimental heat maps, (B) and (D) shows the simulated results. (E-H) show the distributions (purple for simulation and blue for the experimental data) of the RF relative position along the $x$-axis (E), velocity (F), peak velocity values (G) and the consecutive time between velocity peaks (H). We used the same parameters as in Fig.2. For each VF velocity, we ran 100 simulations (in which the RF initial position was random) each for 5000$[s]$ (500,000 iteration steps).

### S12. RF FOLLOWING 2VF

We present here extended data for the case of the RF following 2VF (complimentary to Fig.3).

Note that we could postulate that the fish makes weaker burst forces ($f_0$) when the frequency of its tail bursts decreases, which could improve the agreement, but we opted to keep the model as simple as possible, and maintain that all the parameters are independent of the average VF speed. In particular, we chose a burst force amplitude that would prevent the RF from losing the fastest moving VF (0.07$[m/s]$), which causes it to move too close to the VF for the slower moving VF cases.

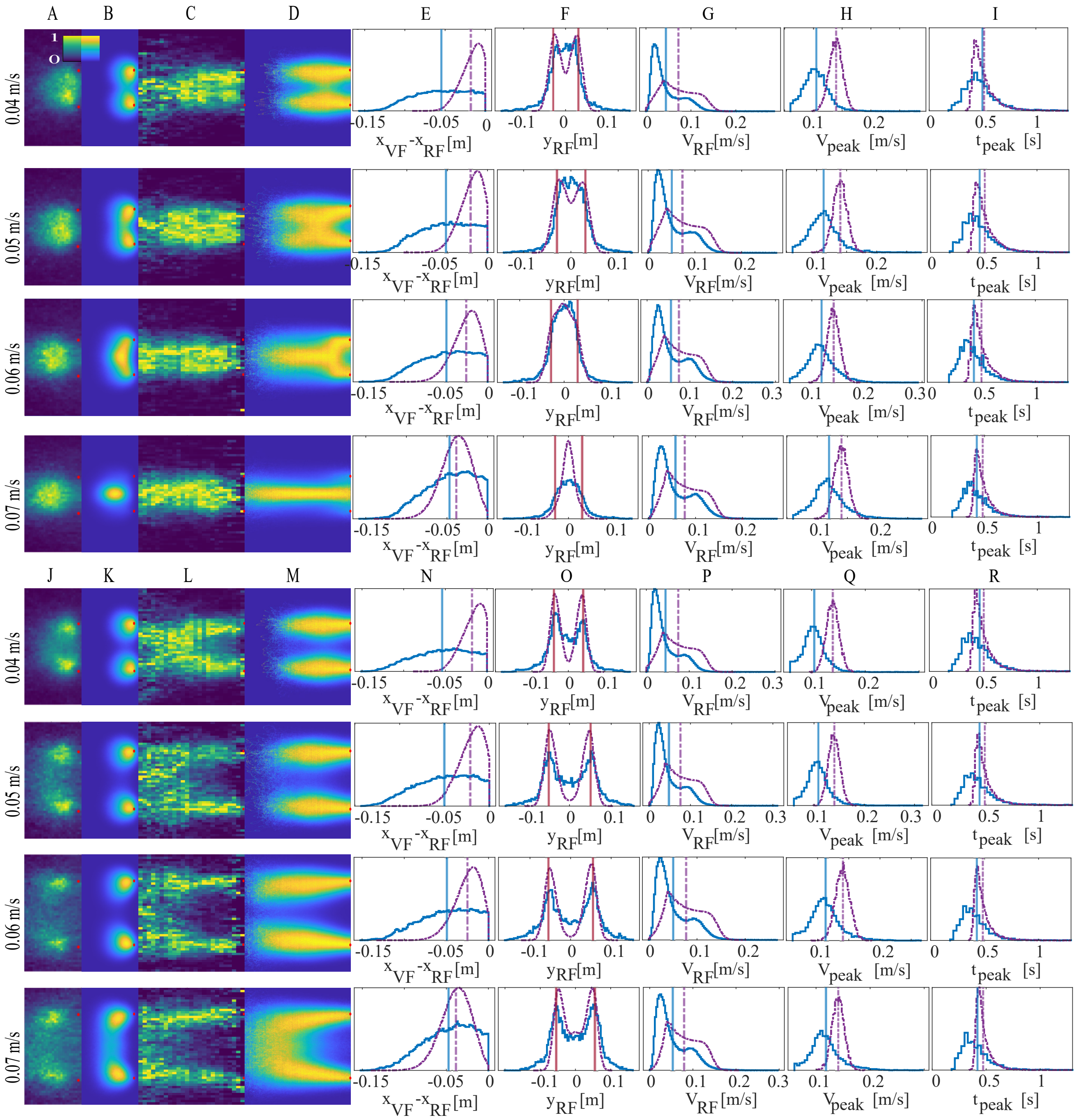


FIG. S12: Two VF - heat maps normalized over the whole 2D space (A-B and J-K) or over the $x$-axis sections (C-D and L-M). (A,J) and (C, L) The experimental RF's accumulated distribution, while (B, K) and (D, M) show the corresponding simulated distributions. (E, N) Distributions of the RF's $x$-positions relative to the 2VF, and $y$-positions (F, O). (G, P) Histograms of the RF speed ($V_{RF}$), (H) burst velocity peak values ($P_{peak}$), and (I) the time intervals between consecutive velocity peaks ($t_{peak}$). All panels compare the experimental data (blue) with the model (purple). The average of each distribution is given by the vertical line, with the corresponding color. The top four lines are for $LRD = 0.06m$, while the bottom four lines are for: $LRD = 0.08[m]$ for $V_{VF} = 0.04[m/s]$, $LRD = 0.1[m]$ for $V_{VF} = 0.05[m/s]$, and $LRD = 0.11[m]$ for $V_{VF} = 0.06[m/s], 0.07[m/s]$). We used the same parameters as in the 1VF system (Fig.2), except for: $f_0 = 1.2[m/s^2]$. When dealing with more than one VF we also use the following parameters: $\nu = 0.5$ and $\sigma_\theta = 0.00002\pi$. For each VF velocity, we ran 100 simulations (in which the RF initial position was random) each for 5000$[s]$ (500,000 iteration steps).

### S13. RF FOLLOWING 3VF

We present here extended data for the case of the RF following 3VF (complimentary to Fig.4).

The distributions of the RF speed are in reasonable agreement between the simulations and the experiments (Fig.S13F), while the distributions of the velocity peaks and the time intervals between consecutive velocity peaks are in very good agreement (Fig.S13G,H).

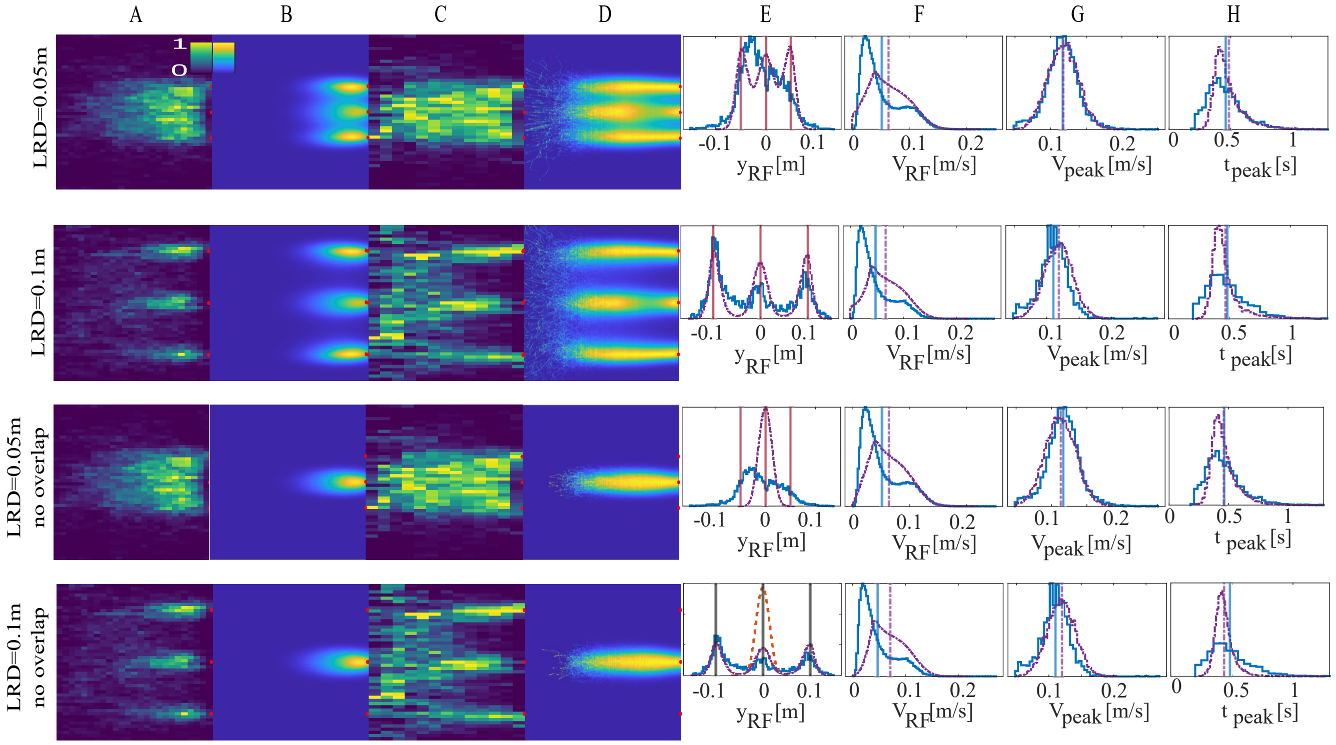


FIG. S13: RF following 3VF. The LRD value for each line in (A-H) is given on the left, with the lowest two lines presenting simulations without the overlap function. Accumulated spatial RF distribution, normalized over the whole 2D space (A,B) or over the $x$-axis sections (C,D). (A,C) Experimental data, while (B,D) the simulation results. (E) Distributions of the RF's projected $y$-positions, (F) RF's speed, (G) Speed of the RF at the peaks, and (H) the time intervals between consecutive speed peaks. In (E-H) we compare the experimental data (blue) with the model (purple). The average of each distribution is denoted by the vertical line, with the corresponding color. In (A-H) the VF velocity is $0.04[m/s]$. We used the same parameters as in Fig.2, except for the burst force amplitude which was changed to $f_0 = 0.95[m/s^2]$. we ran 100 simulations (in which the RF initial position was random) each for $5000[s]$ (500,000 iteration steps).

### S14. RF FOLLOWING 2VF IN SHIFTED GEOMETRY

We present here extended data for the case of the RF following 2VF in shifted geometry (complimentary to Fig.5).

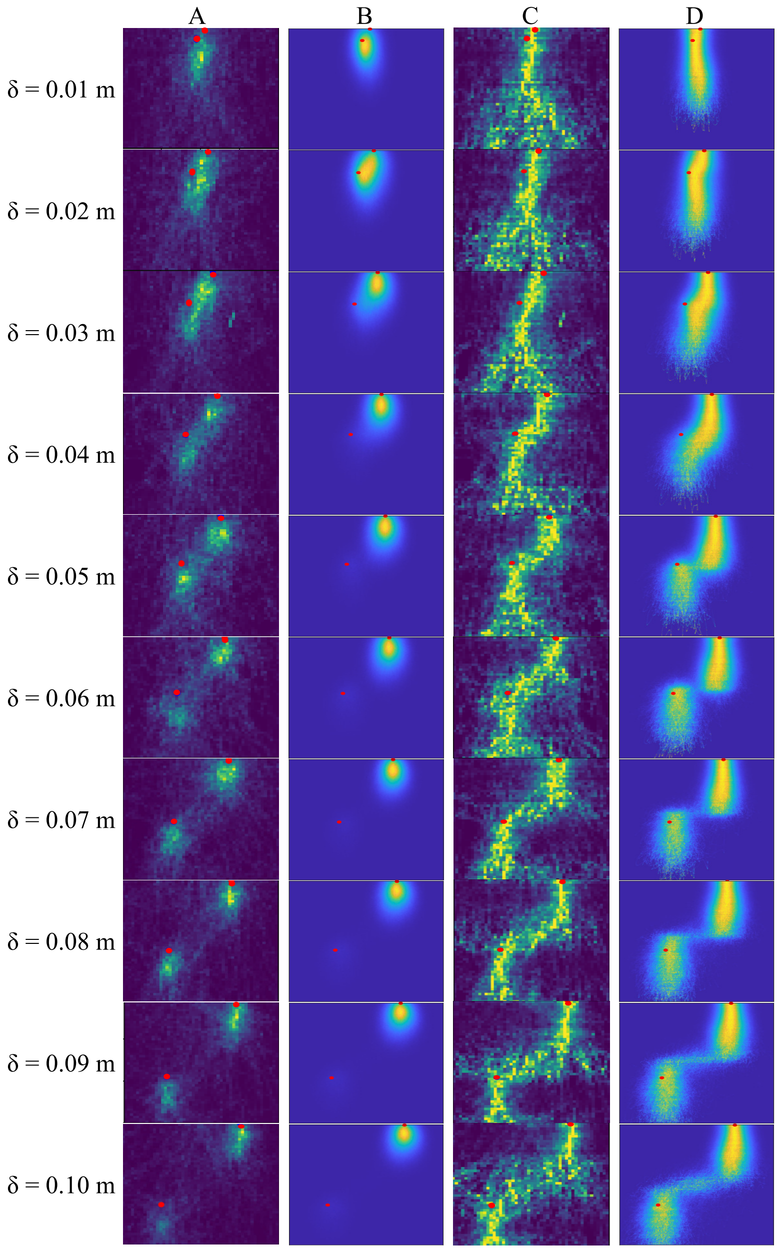


FIG. S14: 2VF in a shifted geometry: one of the VF is shifted in the front-back and the left-right directions with respect to the other VF, by $\delta$. The accumulated spatial distribution of the RF behind the 2VF, (A,B) normalized over the whole 2D space, or (C,D) normalized over $x$-axis sections. (A) and (C) show the results from experimental data, while (B) and (D) show results from the simulations. The VF are denoted by the red circles. The model parameters are the same as used in Fig.S12. The right panels show examples of RF trajectories relative to the 2VF (red dots), for the cases of $\delta = 0.03[m]$, and $\delta = 0.06[m]$. In both plots the color of the line represent the time $[s]$.

### S15. MANY RF

We may now ask what happens when a group of RF interact with each other according to our model. In Fig.S15A,B we plot examples of typical trajectories for either 2RF or 3RF, in the presence of a single persistent leader, in the form of a single VF that moves along a circle (green, see also Supplementary Movies M7-M9). The RF start at the positions indicated by the full circles. We find that at an early time the RF follow the VF, as indicated by the "Following" state (at the locations indicated by the stars). At the corresponding times the histograms on the right show the proportion of "on" spins in each RF, indicating that the blue RF has its maximal attention on the VF leader, while the orange RF is following the blue RF. This continues for a certain time, until their attention changes to following each other.

This occurs at the time indicated by the fish shapes on the trajectory, and the corresponding histograms shown at this "separation" time. The RF lose the leader, and form a non-polar shoal. The time duration until this separation event occurs decreases with increasing number of RF, as shown in Fig.S15C (the full distributions of the separation times are given in Fig.S16).

In Fig.S15D we demonstrate that in the absence of any leader fish, our model naturally gives rise to cohesive shoal behavior, without any spontaneous alignment of the fish along a particular direction (see Supplementary Movie M10). This falls into the category of "swarm" behavior, with low global polarization of the velocity vectors [2]. Note that our model ignores the finite size of the RF, which means that we do not describe the short-range maneuvers that are necessary for collision avoidance. Similarly, we ignore obstructions of one RF by another, with all the RF visible at all times. In addition, our model explicitly involves many-body interactions, that are not simply the superposition of pairwise interactions. Significant interactions beyond pairwise have indeed been measured inside fish schools [3].

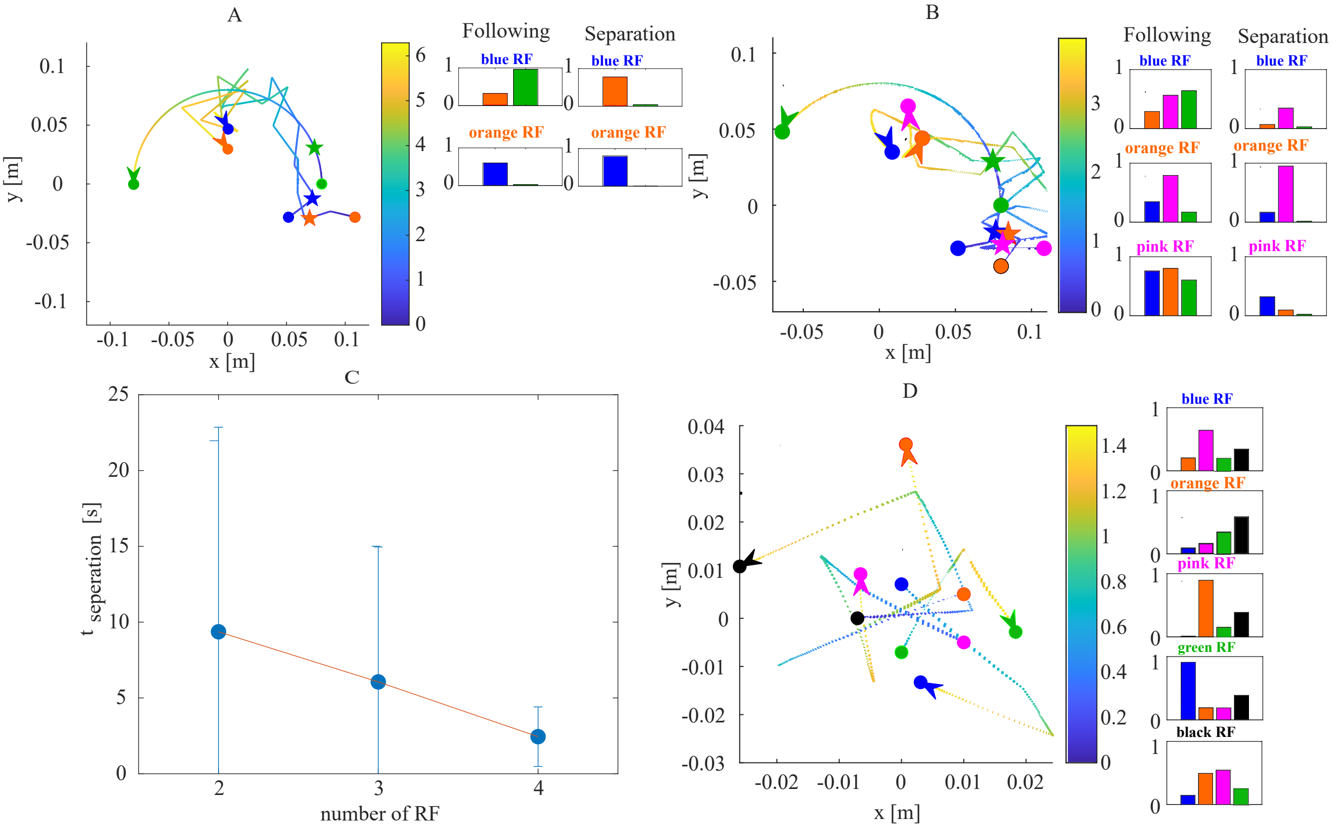


FIG. S15: Simulating groups of RF. (A, B) Examples of simulated trajectories, with (A) 2RF and (B) 3RF in the presence of a VF (green) swimming in a circle of radius $0.08[m]$ with a constant velocity of $0.05[m/s]$. The starting positions of the RF and VF are denoted by the colored circles, and the end points by the colored fish. The side panels show the spin states of each of the RF, representing the weight of the fellow RF and the VF on their direction of motion. We denote these spin states at a typical "following state" (denoted by the stars on the trajectories) where the RF were following the VF, and the "separation state" at the end of the trajectories, where the RF formed a shoal and stopped following the VF. (C) A plot of the mean time until the RF separate from the VF, as a function of the number of RF, based upon running 90,000 simulations for each case of RF number. The mean times for 2RF,3RF and 4RF are: $t(2RF) = 9.36[s] \pm 13.49[s]$, $t(3RF) = 6.05[s] \pm 8.89[s]$ and $t(4RF) = 2.44[s] \pm 1.96[s]$. (D) A typical short trajectory of five RF swimming together with no leader (colored circles denote the starting points). We used the same parameters as in Fig.2.

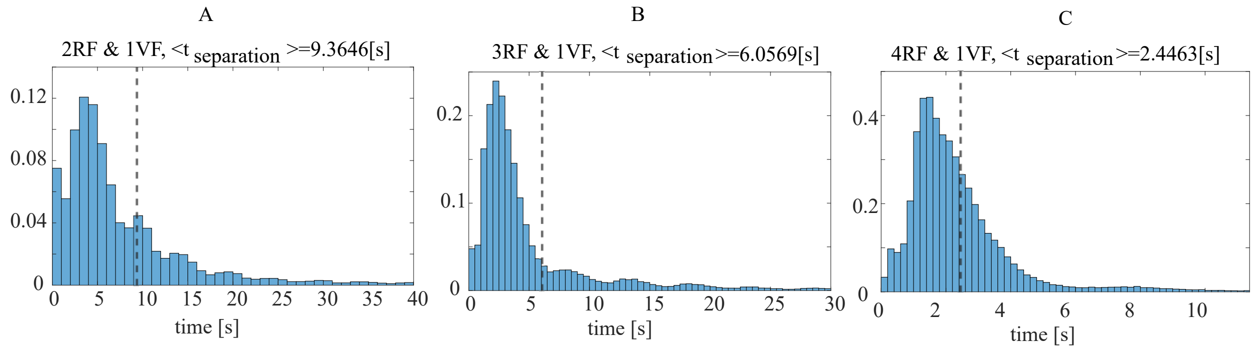


FIG. S16: Separation time histograms for many RF and one VF - the plots show the time of separation for 2RF (A), 3RF (B) and 4RF (C) that are following one VF in a circle of radius $0.08[m]$ with a linear velocity $V_{VF} = 0.05[m/s]$ (Fig.6A,B). The average separation times (Fig.6C) were calculated from conducting $90,000$ simulations. All simulations ran with the same parameters as in the 1VF case (Fig.2).

### S16. SUPPLEMENTARY MOVIES

The simulated RF dynamics while following 1VF, 2VF, and 3VF are shown in movies M1-M6. The many RF following 1VF target moving along a circle are shown in movies M7 (2RF), M8 (3RF), and M9 (4RF). Also, we show the many RF dynamics with no leader among them in movie M10 (5RF). All movies are in the following link: Movies

- Movie M1- one RF following one VF ($V_{VF} = 0.06[\frac{m}{s}]$), the top plot shows the RF in the real frame of reference, and the bottom plot, the RF is simulated relative to the VF (which is in the origin). The RF is in blue and the VF is in red in both cases (parameters as in Fig.2).
- Movie M2 - one RF (blue) following 2VF (red) in the compromise regime ($LRD = 0.06[m]$, and $V_{VF} = 0.07[\frac{m}{s}]$), the right plot shows the neuronal firing in the RF brain, in which "1" represent neurons directed at the VF in $y = 0.03[m]$, and "2" represent neurons directed at the VF in $y = -0.03[m]$ (parameters as in Fig.3).
- Movie M3- one RF (blue) following 2VF (red) in the bifurcated regime ($LRD = 0.11[m]$, and $V_{VF} = 0.07[\frac{m}{s}]$), the right plot shows the neuronal firing in the RF brain, in which "1" represent neurons directed at the VF in $y = 0.055[m]$, and "2" represent neurons directed at the VF in $y = -0.055[m]$ (parameters as in Fig.3).
- Movie M4- one RF (blue) following 2VF (red) in the shifted configuration ($LRD = 0.03[m]$, $V_{VF} = 0.04[\frac{m}{s}]$, and $\delta = 0.03[m]$), the right plot shows the neuronal firing in the RF brain, in which "1" represent neurons directed at the VF in $y = 0.03[m]$, and "2" represent neurons directed at the VF in $y = -0.03[m]$ (parameters as in Fig.3).
- Movie M5- one RF (blue) following 3VF (red) in the compromise regime ($LRD = 0.05[m]$, and $V_{VF} = 0.04[\frac{m}{s}]$), the right plot shows the neuronal firing in the RF brain, in which "1" represent neurons directed at the VF in $y = 0.05[m]$, "2" represent neurons directed at the VF in $y = 0[m]$, and "3" represent neurons directed at the VF in $y = -0.05[m]$ (parameters as in Fig.4).
- Movie M6- one RF (blue) following 3VF (red) in the bifurcated regime ($LRD = 0.10[m]$, and $V_{VF} = 0.04[\frac{m}{s}]$), the plot on the right side shows the neuronal firing in the RF brain, in which "1" represent neurons directed at the VF in $y = 0.1[m]$, "2" represent neurons directed at the VF in $y = 0[m]$, and "3" represent neurons directed at the VF in $y = -0.1[m]$ (parameters as in Fig.4).
- Movie M7- 2RF following 1VF (green) in a circle (of radius $0.08[m]$). The neuronal activity for each RF are given in the right bar plots, where the bar color indicates a neuronal group directed at the fish with the same color (parameters as in Fig.2).
- Movie M8- 3RF following 1VF (green) in a circle (of radius $0.08[m]$). The neuronal activity for each RF are given in the right bar plots, where the bar color indicates a neuronal group directed at the fish with the same color (parameters as in Fig.2).
- Movie M9- 4RF following 1VF (green) in a circle (of radius $0.08[m]$). The neuronal activity for each RF are given in the right bar plots, where the bar color indicates a neuronal group directed at the fish with the same color (parameters as in Fig.2).

- Movie M10- Simulation of 5RF (a shoal) with no leader. The right plot shows the polarization vector- the average direction of the whole shoal (parameters as in Fig.2).


[1] V. H. Sridhar, L. Li, D. Gorbonos, M. Nagy, B. R. Schell, T. Sorochkin, N. S. Gov, and I. D. Couzin, Proceedings of the National Academy of Sciences **118** (2021).
[2] J. Delcourt and P. Poncin, Reviews in Fish Biology and Fisheries **22**, 595 (2012).
[3] Y. Katz, K. Tunstrøm, C. C. Ioannou, C. Huepe, and I. D. Couzin, Proceedings of the National Academy of Sciences **108**, 18720 (2011).
